# From Monomers to Oligomers: Structural Mechanism of Receptor-Triggered MyD88 Assembly in Innate Immune Signaling

**DOI:** 10.1101/2024.09.13.612588

**Authors:** Kazuki Kasai, Kayo Imamura, Masatoshi Uno, Naotaka Sekiyama, Tomoko Miyata, Fumiaki Makino, Ryusei Yamada, Yoshiki Takahashi, Noriyuki Kodera, Keiichi Namba, Hidenori Ohnishi, Akihiro Narita, Hiroki Konno, Hidehito Tochio

## Abstract

MyD88 plays a pivotal role in Toll-like receptor (TLR) and interleukin-1 family signaling through its oligomerization upon receptor activation, leading to downstream protein recruitment. The Toll/interleukin-1 receptor domain of MyD88 (TIR_MyD88_) is responsible for this receptor-mediated oligomerization, but the detailed mechanism involved remains elusive. We investigated the structure of TIR_MyD88_ oligomers and their interactions with TLRs. Cryoelectron microscopy revealed that tandemly arrayed TIR_MyD88_ subunits formed an antiparallel double-stranded filament that could further form rings and cylindrical filaments. Moreover, the self-assembly of TIR_MyD88_ *in vitro* was markedly accelerated by dimeric rather than monomeric receptor TIRs, possibly reflecting the signal initiation step *in vivo*. High-speed atomic force microscopy further captured the dynamic processes of oligomerization of TIR_MyD88_, in addition to its direct interaction with the receptor TIRs. Based on these results, a novel regulatory mechanism of TIR_MyD88_ oligomerization underlying the signal initiation step was revealed.

## Introduction

Myeloid differentiation primary response 88 (MyD88) is the main cytosolic adaptor protein involved in innate immune and inflammatory signaling ^1,2^. It interacts with interleukin-1 receptor (IL-1R) family members and most members of the Toll-like receptor (TLR) family. Upon ligand binding, these receptors interact with MyD88, which subsequently recruits and activates members of the IL-1R-associated kinase (IRAK) family, such as IRAK4 and IRAK1.

MyD88 consists of two protein‒protein interaction domains: the death domain (DD_MyD88_) at the N-terminus and the Toll/IL-1R domain (TIR_MyD88_) at the C-terminus. These domains are connected by a linker of 46 amino acid residues called the intermediate domain (ID) ^3,4^. Studies have shown that MyD88 has intramolecular interactions between DD_MyD88_ and TIR_MyD88_, which may play a regulatory role in signal transduction ^5,6^. On the other hand, in general, both the DD and TIR domains participate in homotypic interactions, leading to the formation of oligomeric signaling complexes ^2,7^. During IL-1R and TLR signaling, DD_MyD88_ forms helical homo-oligomers ^8^ that serve as scaffolds for the successive recruitment of IRAK4 and IRAK1/2. The details of the interaction governing this recruitment were clarified by the determination of the tripartite complex structure of DD_MyD88_/DD_IRAK4_/DD_IRAK2_ ^9^, in which a three-layer DD oligomer is formed by the respective DDs at a 6:4:4 ratio. It is speculated that the formation of this DD oligomer brings the kinase domains of the IRAKs into close proximity, facilitating transphosphorylation, which triggers their activation and the subsequent phosphorylation of downstream proteins, such as TRAF6 ^2^.

While DD_MyD88_ is responsible for recruiting and activating IRAKs, TIR_MyD88_ plays a crucial role in interacting with IL-1Rs/TLRs through homotypic TIR-TIR interactions. Signaling by these receptors is initiated by ligand-induced dimerization of their extracellular domains, which leads to subsequent dimerization of their cytosolic TIR domains. The dimerized receptor TIRs then engage with TIR_MyD88_, triggering the formation of MyD88 clusters at the receptor site ^10^. In some but not all TLR signaling modes, another adaptor protein, MyD88 adaptor-like protein (Mal) plays an essential role by mediating interactions between TLRs and MyD88 ^11,12^. A recent live-cell imaging study revealed that in IL-1 signaling, the formation of MyD88 clusters of a certain size is necessary for the recruitment of IRAK4 and the initiation of appropriate signaling. Thus, the authors proposed that such clustering functions as a “physical threshold” for signaling ^13^.

Although monomeric structure of TIR_MyD88_ was reported years ago ^14,15^, the manner in which TIR_MyD88_ oligomerizes has remained elusive. Recently, Ve et al. reported that TIR_MyD88_ forms a fibrous structure in the presence of the TIR domain of Mal (TIR_Mal_) ^16^. Clabbers et al. subsequently grew crystals of TIR_MyD88_ by seeding with filaments of TIR_Mal_ ^17^, demonstrating that in the crystals, the TIR_MyD88_ subunits were arrayed in a manner identical to that observed for the subunits in the seeded TIR_Mal_ filaments. Therefore, the authors propose a molecular templating mechanism, in which TIR_Mal_ assemblies act as templates for the assembly of TIR_MyD88_. Although such templating may be a role for TIR_Mal_, it is important to note that Mal is not required for many of the MyD88-dependent signals. For instance, while Mal is essential for a subset of TLRs (such as TLR2 and TLR4) ^2,12^, it is not required for signaling through other TLRs, such as TLR5 and TLR7. Even TLR2 can initiate MyD88-dependent signaling without Mal ^18^. Furthermore, the IL-1 family (IL-1α, IL-1β, IL-18 and IL-33) does not require Mal for MyD88-dependent signaling ^19,20^. In such cases, Mal-templated oligomerization cannot occur and thus the receptors directly trigger the self-assembly of TIR_MyD88_. Therefore, for a comprehensive understanding of MyD88-depenedent signaling, it is crucial to elucidate the self-assembly mode of TIR_MyD88_ in the absence of Mal. Furthermore, the detailed molecular process by which the receptors trigger TIR_MyD88_ assembly remains elusive. In addition to these mechanistic interests, the self-assembly mode of TIR_MyD88_ is of clinical importance. In aggressive subclasses of B-cell lymphoma, somatic gain-of-function mutations in TIR_MyD88_, such as L265P (also referred to as L252P), are frequently identified. These mutations lead to an unusually pronounced self-assembly of MyD88 independent of receptors as well as Mal, resulting in aberrant activation of the NF-κB and MAP kinase pathways. This unregulated activation is a primary cause of increased tumor malignancy ^21–23^. Additionally, these mutations are implicated in autoinflammatory diseases and severe arthritis ^24,25^. Therefore, the molecular details of the TIR_MyD88_ self-assembly process may also provide valuable insights into the pathological mechanisms underlying these diseases and facilitate the development of new therapeutics.

Here, we show that even in the absence of Mal, TIR_MyD88_ self-assembles into filaments that can further form ring structures and cylindrical fibers. Notably, the addition of a small amount of dimeric receptor TIRs markedly promotes the self-assembly of TIR_MyD88_, consistent with the physiological role of receptor dimerization *in vivo*. Through cryoelectron microscopy (cryo-EM), we reveal the self-assembled structure of TIR_MyD88_, which is globally different from the previous Mal-induced TIR_MyD88_ oligomers, albeit with partial similarity, providing vital functional insights. We further visualize the dynamics of the TIR_MyD88_ oligomerization and capture its direct binding to receptor TIRs via high-speed atomic force microscopy (HS-AFM), demonstrating that a specific loop reorganization acts as a kinetic barrier for oligomerization of TIR_MyD88_. Combining these results with those of structure-guided cell-based assays, we propose a unique regulatory mechanism for the receptor-triggered assembly of MyD88, providing molecular insights into the formation of Myddosome signaling complex ^9^ and related pathogenetic mechanisms.

## Results

### Self-assembled oligomers of TIR_MyD88_

We expressed and purified a recombinant TIR domain of MyD88 (TIR_MyD88_; residues 172–309) and found that it spontaneously assembled into filaments when incubated in purification buffer (10–300 μM). The filaments also formed rings with a characteristic diameter of 22-38 nm (Fig. 1a, left) and, occasionally, larger rings with a diameter of 60-80 nm. Moreover, as the incubation time increased, short cylindrical fibers were formed (Fig. 1a, middle), resulting in a predominance of long, straight cylindrical fibers (Fig. 1a, right). Although the global morphology varied with incubation time (rings or cylinders), it appeared that all morphologies were formed from the common filaments observed during early incubation. This conclusion was based on the observation that such filaments projected from disordered/degraded cylindrical structures (Fig. 1b). Thus, we considered the filaments to be fundamental for the formation of the rings and cylindrical fibers. To characterize the time course of TIR_MyD88_ self-assembly, turbidity assays were conducted, and the results showed a sharp increase in the assembly rate for TIR_MyD88_ concentrations greater than around 200 μM (Fig. 1c,d). Whereas we found that TIR_MyD88_ formed fibers at concentrations of 10 μM or above, it was previously reported that it had no tendency to form fibrils by itself ^16^. One possible explanation for this inconsistency is the presence of extra amino acid residues, including a purification tag (His-tag) at the C-terminus of TIR_MyD88_ in the previous study. These additional residues may have perturbed the intricate intermolecular interactions, preventing TIR_MyD88_ self-assembly.

**Fig. 1:**
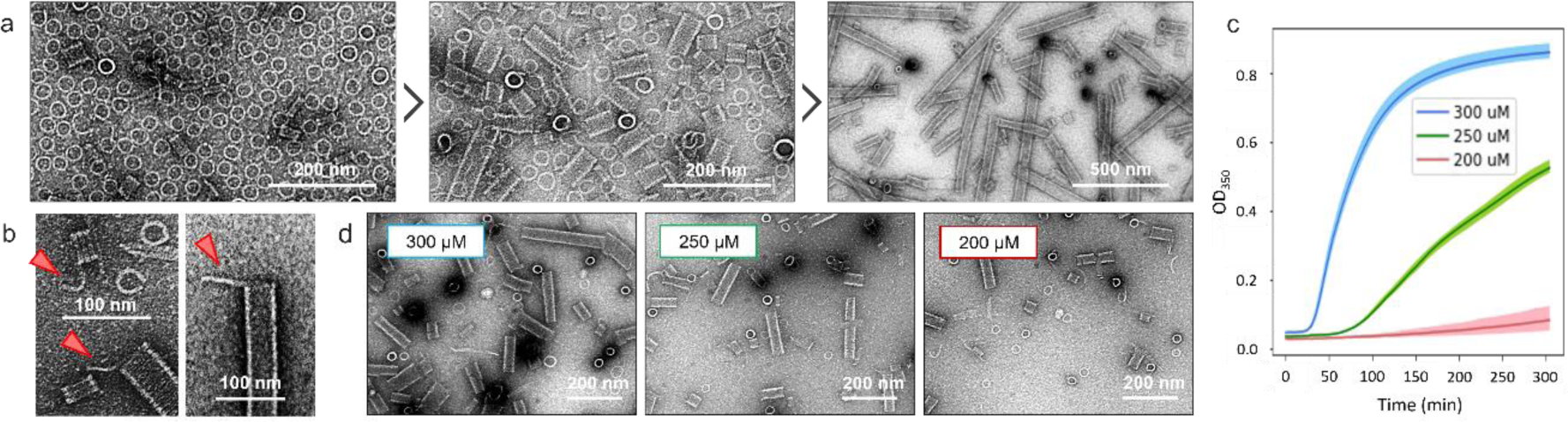
Time course of TIR_MyD88_ self-assembly. (a) Negative stain TEM images of TIR_MyD88_ (200 μM) recorded after different incubation times (2 hours, 1 day, and 3 days) at 30°C. (b) The free filaments projected from the cylindrical fibers are indicated by red triangles. (c) A turbidity assay was performed in triplicate at 37°C with different concentrations of TIR_MyD88_. The optical density at 350 nm (OD_350_) was measured every 5 min. (d) TEM images of fiber samples collected after the turbidity assay described in (c).

### HS-AFM reveals that TIR_MyD88_ can form ring structures

To determine whether TIR_MyD88_ forms filaments, we employed HS-AFM ^26,27^, which enables single-molecule visualization of biomacromolecules in physiologically relevant aqueous solutions. Although only a sparse distribution of monomeric TIR_MyD88_ was observed in diluted solution (10 nM) ^5^, incubation with higher concentrations (10-200 μM) resulted in the formation of ring structures, with sizes and shapes similar to those observed by TEM, within 10 min (Fig. 2b,c). The height and diameter of the rings were 6-7 nm and 24-36 nm (average, 29 nm), respectively (Fig. 2 and Extended Data Fig. 1). Examination of the nodules on the surface of one typical ring led to the identification of 29 subunits (each ∼3 nm in diameter) on the surface (Fig. 2a,b). Because the height of the ring corresponded to that of two monomers of TIR_MyD88_, the ring was anticipated to be double layered, indicating that it contained a total of 58 subunits (Fig. 2c). The double-layered structure was evident in cross sections of a partially formed ring. The rings contained two distinct regions with different heights, i.e., 6.5 nm and 3.2 nm (Extended Data Fig. 1), corresponding to double- and single-layered regions, respectively.

**Fig. 2:**
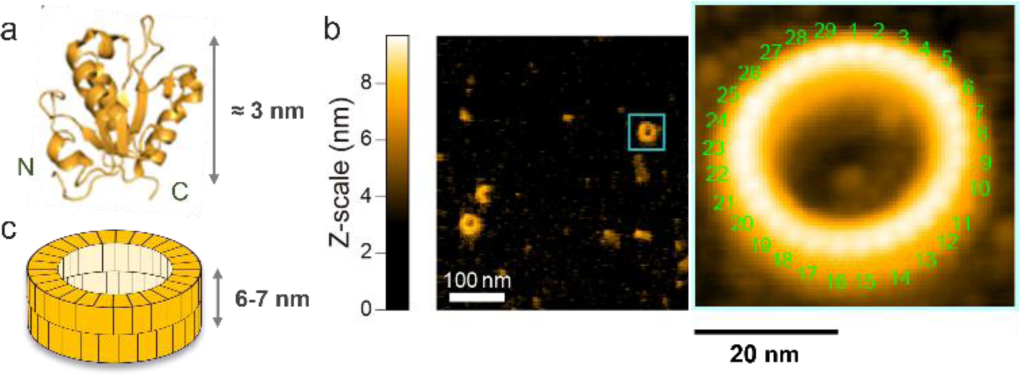
AFM images of TIR_MyD88_ rings. (a) Ribbon model of monomeric TIR_MyD88_ (PDB ID: 2Z5V). (b) Left: representative overview of the rings as visualized by AFM. Scan area: 500 × 500 nm^2^ with 150 × 150 pixels. Scan speed: 600 ms per image. Right: magnified view of the ring enclosed in the cyan square in (b). Scan area: 60 × 60 nm^2^ with 120 × 120 pixels. Scan speed: 600 ms per image. (c) Schematic of the ring shown in (b).

Notably, we did not observe cylindrical fibers by AFM, as we did by TEM (Fig. 1). This difference may be due partially to the short incubation time of TIR_MyD88_ on the AFM stage before measurement (5∼10 min). In addition, in AFM, mechanical forces are applied to the specimen; thus, only molecular assemblies that are somewhat resistant to deformation are visualized. The interactions that stabilize the cylindrical fibers may be weaker than those that form the double-layered rings.

### Determination of the structure of the cylindrical fibers of TIR_MyD88_ by cryo-EM

The ability of TIR_MyD88_ to form filaments and higher-order assemblies under physiological conditions suggests that the observed self-assembly mode represents a functional state of TIR_MyD88_ in cells. Furthermore, gain-of-function mutations identified in TIR_MyD88_ have been reported to be relevant to refractory B-cell lymphomas, such as activated B-cell-like diffuse large B-cell lymphoma (ABC-DLBCL) ^21^, as well as a subset of Schnitzler’s syndrome cases ^24^. These mutations, particularly L265P (L265 is referred to as L252 in this study), have been shown to result in aberrant self-assembly of TIR_MyD88_ and constitutively activate downstream signals that promote tumor survival ^21,22^. Given the anticipated functional and clinical significance of the self-assembled structure of TIR_MyD88_, we sought to determine the structure of the cylindrical fibers using cryo-EM.

The obtained micrographs showed cylindrical fibers with diverse widths. Two types of frequently observed cylinders were selected for analysis: thinner cylinders (∼28 nm diameter) and thicker cylinders (∼36 nm diameter) (Extended Data Fig. 2a and 5a). We first analyzed the thinner cylinders. After 2D class averaging, the images clearly showed double-layered rings stacked coaxially (Fig. 3a). This structure was reminiscent of the double-layered rings observed by AFM. In fact, the height of each double ring in the cylinder was 6.6 nm, which was consistent with that of the rings observed by AFM (Fig. 2 and Extended Data Fig. 1). Next, 3D helical reconstruction was performed on the 26,680 selected images, resulting in a 3D EM map with a resolution of 3.3 Å (FSC 0.143), and an atomistic model was constructed (Fig. 3b,c, Extended Data Fig. 2 and Supplementary Table 1).

**Fig. 3:**
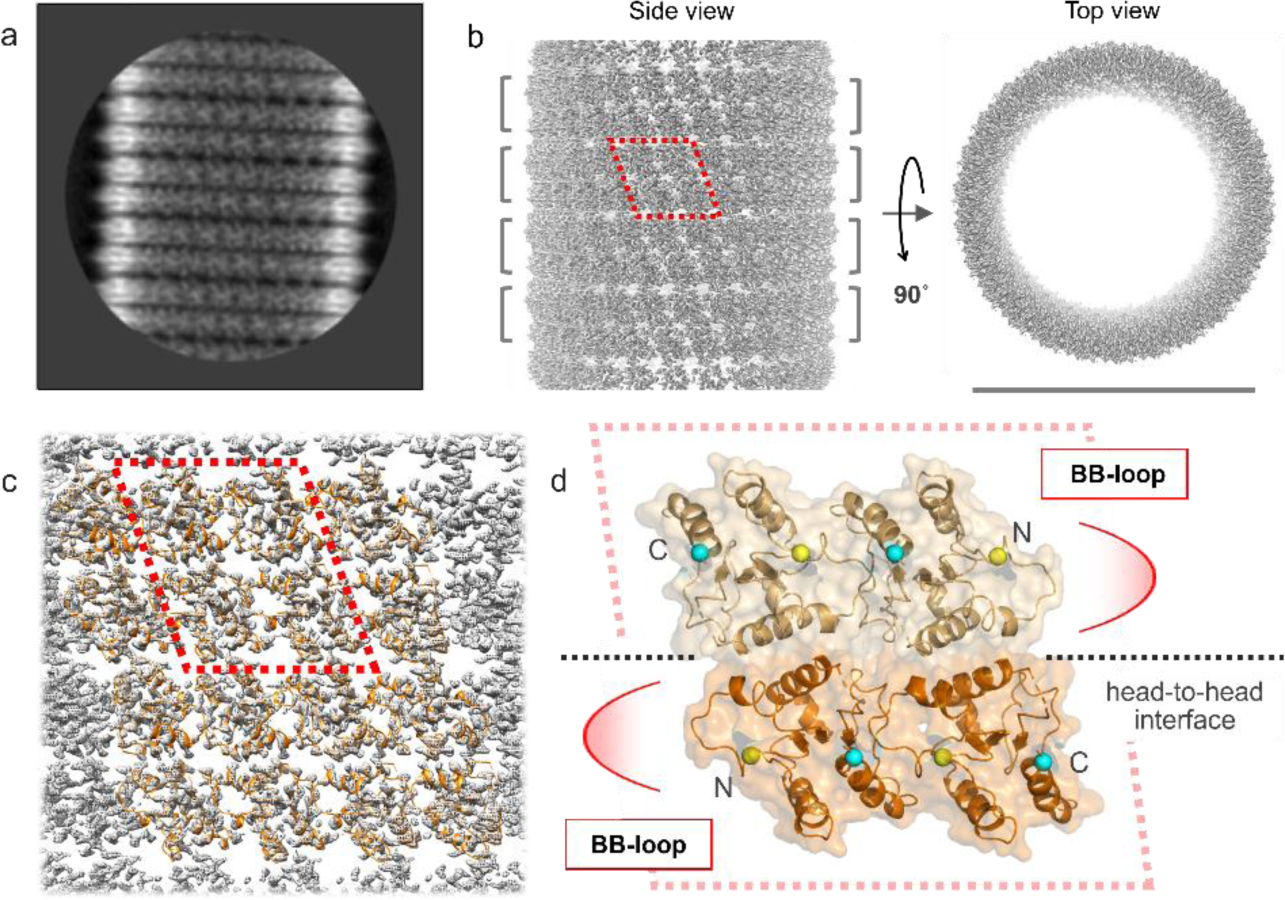
Cryo-EM structure of the cylindrical fibers of TIR_MyD88_. (a) Representative image of a cylindrical fiber of self-assembled TIR_MyD88_ after 2D class averaging. (b) Cryo-EM density map of the cylindrical fiber in two orientations. The gray brackets indicate the positions of the double-layered rings. The gray bar under the top view represents a length of 28 nm. (c) Molecular models fitted to the EM map. (d) Depiction of the four subunits enclosed in the red dotted outlines in (b) and (c) in ribbon and surface representations. The red arcs and black dotted lines indicate the intrastrand and head-to-head stacking interfaces, respectively. The N- and C-termini are depicted as yellow and cyan spheres, respectively.

### Structural characterization of the filaments and cylinders

Overall, each cylindrical fiber consists of coaxially stacked double-layered rings ∼28 nm in diameter, each containing 26 TIR_MyD88_ subunits per layer in each round (Fig. 3a,b). Each of the double-layered rings consists of two identical rings stacked head-to-head (Fig. 3c,d). To form the cylinder, the double-layered rings are stacked coaxially tail-to-tail. The head-to-head stacking interface (referred to as “the dense interface”) is more densely packed than the tail-to-tail interface (“the sparse interface”); the buried surface areas of the dense and sparse interfaces are ∼440 Å^2^ and ∼337 Å^2^ per subunit, respectively. Detailed examination of the structure revealed that the subunit arrangement in the filament (i.e., equivalent to the double ring) is partially identical to that in the two-stranded filament of TIR_MyD88_ crystallized with the aid of Mal filaments ^17^. However, there are considerable differences between our filaments and the Mal-induced oligomers. In particular, the alignment of the two strands is completely opposite. That is, in the Mal-induced oligomers, two TIR_MyD88_ strands are joined in a parallel orientation, whereas in our filaments, the two strands are in an antiparallel orientation (Fig. 3d and Extended Data Fig. 3a-c). Furthermore, in the Mal-induced oligomers, each subunit in one of the strands is wedged between two adjacent subunits in the other strand, whereas in our cryo-EM structure, one subunit in one strand is in complementary and symmetrical contact with one subunit in the other strand. In a sense, this *antiparallel double-stranded* filament can be considered a linear assembly composed of symmetric head-to-head dimers of TIR_MyD88_ (Fig. 3d and Extended Data Fig.3a).

For convenience, we can categorize the subunit interactions in the cylinders into three types: “intrastrand,” describing interactions within individual strands; “interstrand”, describing interactions at the dense interface; and “interfilament”, describing interactions at the sparse interface (Fig. 4a). Intrastrand interactions account for 8∼9% (∼745 Å^2^) of the total surface area of a single subunit. These interactions are essentially identical to those in the Mal-induced oligomers. Based on previously proposed nomenclature ^17^, the BB surface of one subunit binds to the EE surface of the adjacent subunit (Fig. 4a and Extended Data Fig. 2c), and the protein backbone structure in these regions is similar to that in the Mal-induced oligomers (Extended Data Fig. 3d-i). On the other hand, dramatic differences from the Mal-induced oligomers were evident in the interstrand interactions, accounting for ∼5% (∼440 Å^2^) of the total surface area of a single subunit, in which the αB and αC helices (the BC surface) from two subunits interact with each other (Fig. 4a). In a sense, the two subunits form a head-to-head dimer with C2 symmetry through the BC surface residues, in which W205, F235, K238, F239, and S242 are positioned symmetrically at the interface to participate in extensive hydrophobic interactions with each other (Fig. 4a and Extended Data Fig. 3j). The resultant hydrophobic core is supported by peripheral hydrogen bonds between W205 and S242. The symmetric dimers assemble linearly through intrastrand interactions to form a double-stranded filament, in which diagonal interactions also play a role, with hydrophobic contacts between I267 and F270 and hydrogen bonds between S266 and R269 (Fig. 4a). Consequently, the lateral surface of the filament is markedly distinct from that of the Mal-induced oligomers. A notable difference is the location of the N-termini to which DD_MyD88_ should be connected. In our double-stranded filaments (i.e., the rings), all the N-termini are on one side of the filament, whereas in the Mal-induced oligomers, they are directed toward the opposite sides of the oligomers (Extended Data Fig. 3a,b). DD_MyD88_ forms a helical oligomer that serves as the acceptor site for IRAK4. The subunit arrangement in our double-stranded filaments is expected to facilitate the formation of this oligomer because four TIR_MyD88_ subunits form a parallelogram with all four N-terminal ends facing in one direction (red dotted lines in Fig. 3b-d). This arrangement is suitable for the alignment of four DD_MyD88_ subunits to form a helical oligomer because in the oligomer, the C-terminal ends of the four DD_MyD88_ subunits are in a quasisquare configuration (Extended Data Fig. 4). Thus, the formation of the TIR_MyD88_ filament is expected to promote DD_MyD88_ oligomerization (see the Discussion for further details).

**Fig. 4:**
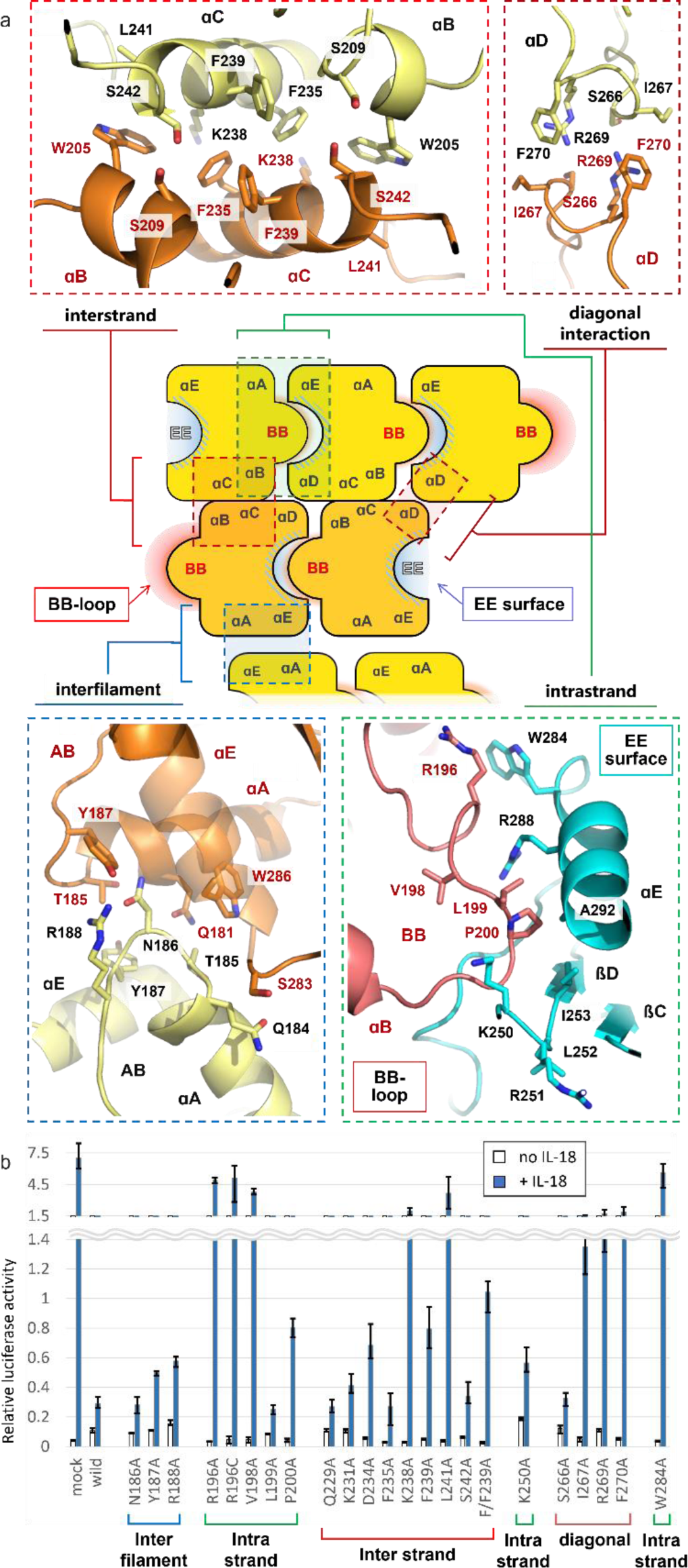
Validation of the double-stranded filament structure. (a) Cartoon showing the arrangement of subunits in the cylindrical fiber and the interfaces of the interstrand (red dotted square), diagonal (dark red dotted square), interfilament (blue dotted square) and intrastrand (green dotted square) interactions depicted with stick models on the ribbon diagrams. (b) NF-κB reporter assay with IL-18-stimulated 293T cells. Each column shows the NF-κB-driven luciferase activity of stimulated (blue) and unstimulated (white) cells expressing WT TIR_MyD88_ or mutants of TIR_MyD88_. F/F239A: double mutant, F235A and F239A. Data shown are the mean and range of triplicate experiments.

The interfilament interactions (the sparse interface) result in the formation of cylindrical fibers from double-layered rings, in which residues from the end of the αA to the AB loop as well as the N-terminal half of the αE play a major role (Fig. 4a). This interface buries 3-4% (∼ 337 A^2^) of the total surface area of a single subunit. This sparse interaction accounts for the observation that a long incubation time is required for the cylindrical fibers to form. Intriguingly, the interfilament interactions, namely, the formation of tail-to-tail dimers, are not symmetric (Fig. 4a). For example, the backbone oxygen of Q181 in one subunit forms hydrogen bonds with the NH side chain of N186 in the other subunit, but the opposite does not occur (i.e., no hydrogen bonds are formed between N186 of the former subunit and Q181 of the latter subunit). Additionally, the OH side chain of Y187 in one subunit forms a hydrogen bond with the NH side chain of R188 in the other subunit, but again, no hydrogen bonds are formed between R188 of the former subunit and Y187 of the latter subunit. In addition, the hydrophobic contacts between the aliphatic regions of the interface residues are asymmetrical. In fact, while four αE residues (K282, S283, W286 and T287) of one subunit are involved in the interface, only two αE residues (S283 and T287) of the other subunit are involved. Thus, the cylinder has polarity.

### Thicker cylinders are formed by spiral winding of the antiparallel double-stranded filaments

We then analyzed the thicker cylindrical fibers. The obtained structure indicated that the cylinder is formed from a single antiparallel double-stranded filament wound into a spiral (Extended Data Fig. 5). However, the subunits are arrayed in essentially the same fashion as in the thinner cylinder, and the intra- and interstrand interactions, as well as the interfilament interactions, are well conserved (Extended Data Fig. 5g,h). Therefore, the two types of cylinders, with different widths, are formed from a common double-stranded filament. The difference is whether the filament closes to form a ring or extends infinitely in a spiral, and this comes from the cumulative effects of slight variations in the placement of each subunit (Extended Data Fig. 5e,f). The origin of the two distinct types of cylindrical fibers from a common double-stranded filament indicates that this self-assembly mode is an inherent property encoded in the TIR_MyD88_ structure, suggesting its biological significance.

### Functional validation of the filament structure

To verify whether our antiparallel double-stranded filament of TIR_MyD88_ reflects its functional state in cells, we conducted a cell-based assay to assess the dominant-negative inhibitory effect of TIR_MyD88_ on NF-κB activity ^14,28,29^. We ectopically expressed either TIR_MyD88_ or its mutants in HEK293T cells and evaluated NF-κB activation induced by IL-18, a member of the IL-1 family. The expression of wild-type TIR_MyD88_ abrogated NF-κB activity (Fig. 4b, wild), indicating that the expression of TIR_MyD88_ inhibited the proper function of endogenous MyD88. Thus, the ectopically expressed TIR_MyD88_, which lacks DD_MyD88_, is thought to co-assemble with endogenous MyD88 and prevents it from forming functional oligomers to recruit IRAK4. Recently, a dominant-negative inhibitory effect exerted by a short splice variant of MyD88 (MyD88S), which lacks the linker region between DD_MyD88_ and TIR_MyD88_, was investigated. The variant was able to interact with full-length MyD88, but functional oligomers were not formed in the cell ^4^. A similar effect may occur here.

In contrast, in cells expressing one of the TIR_MyD88_ mutants harboring mutations at the intrastrand interface (R196A, R196C, V198A, or W284A), marked recovery of NF-κB activity was observed, revealing the critical importance of these residues in forming co-assemblies with endogenous MyD88 (Fig. 4b). In fact, the importance of these intrastrand residues in both IL-1R ^30^ and TLR signaling ^17,31^ has been established. Negative stain TEM revealed that TIR_MyD88_ with the W284A mutation (located on the EE surface) completely failed to self-assemble (Extended Data Fig 6a). Thus, the reduced dominant-negative effects are likely attributable to the reduced self-oligomerization ability of TIR_MyD88_. However, despite the complete loss of self-assembly *in vitro*, the P200A mutant (positioned at the tip of the BB loop) retained some dominant-negative effects. Thus, this mutation may have not completely disrupted co-assembly between mutant TIR_MyD88_ and endogenous MyD88 in the cellular context.

We then examined the interstrand residues that are unique to our double-stranded filaments (red and dark red dotted squares in Fig. 4a). When K238, L241, I267, R269, or F270 was replaced by Ala, significant recovery of NF-κB activity was observed, suggesting that these residues are critical for forming co-assemblies with endogenous MyD88 (Fig. 4b). Interestingly, despite the K238A TIR_MyD88_ mutant did not interfere with endogenous MyD88 in the cells, it was able to form small rings by itself *in vitro* (Extended Data Fig. 6b). The cryo-EM map of the K238A ring shows that the subunit arrangement is different from that of the WT ring and is incompatible with WT TIR_MyD88_ (Extended Data Fig. 6c-e). Thus, the K238A mutant is essentially not expected to stably co-assemble with endogenous MyD88, though the mutant can self-assemble.

Next, we examined the interfilament interface (Fig. 4a, blue dotted square). Replacement of either N186, Y187, or R188 with Ala resulted in only a small reduction in the dominant-negative inhibitory effect, indicating that these residues have limited contributions to forming co-assemblies with endogenous MyD88 in the cells (Fig. 4b). This finding is consistent with a previous report showing that neither N186A nor R188A affected IL-1β-induced NF-κB activity ^30^. TEM revealed that TIR_MyD88_ with the Y187A mutation formed rings similar to those formed by WT TIR_MyD88_ but largely failed to form cylindrical fibers (Extended Data Fig. 6a). Thus, although the interfilament interactions are required for forming the cylindrical assembly *in vitro*, they are dispensable for functional oligomers in cells. Taken together, these data strongly support that the antiparallel double-stranded filament structure represents the functional oligomeric state of TIR_MyD88_ in cells.

### HS-AFM reveals the mode of subunit incorporation during strand elongation

During HS-AFM measurements of TIR_MyD88_ rings, we often observed disintegration and regeneration of the rings. The disintegration was caused partially by the inevitable scanning forces of the AFM cantilever on the specimen. Closer examination of the ring disintegration/regeneration process (Supplementary Video 1) showed that the upper ring disintegrated while the lower ring remained intact. The upper ring subsequently regenerated on top of the lower ring. The most striking feature was that while disintegration of the upper ring occurred in both directions, regeneration occurred only in a counterclockwise direction (Fig. 5a). In other words, the elongation of the upper strand is unidirectional.

**Fig. 5:**
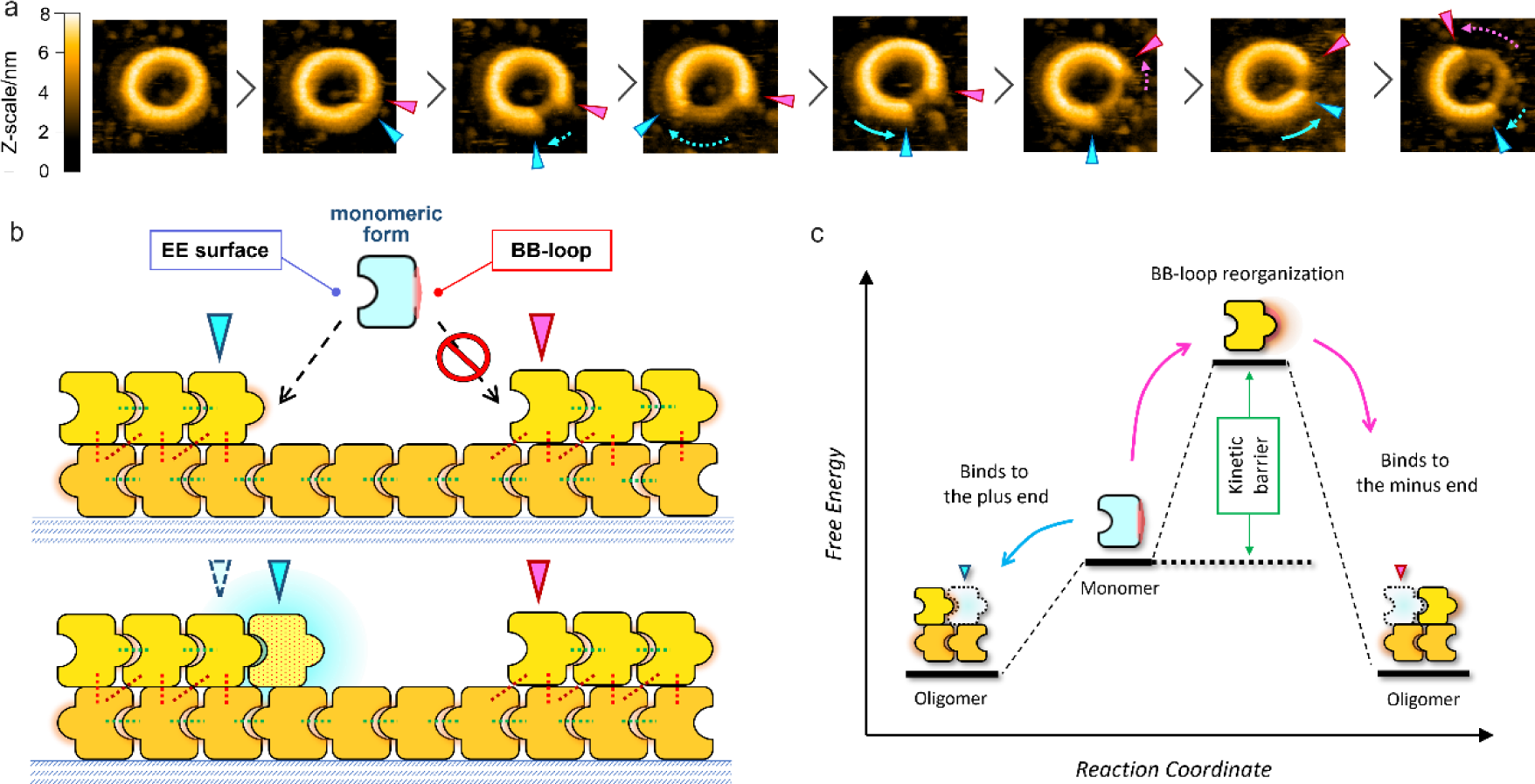
Unidirectional regeneration of the upper ring of TIR_MyD88_. (a) Time series of AFM images showing unidirectional regeneration of the upper ring (Supplementary Video 1). Scan area: 60 × 60 nm^2^ with 120 × 120 pixels. Scan speed: 600 ms per image. Disintegration of the end denoted by the cyan triangle occurs in a clockwise direction (cyan dotted arrow), and regeneration then occurs in a counterclockwise direction (cyan arrow). Disintegration of the other end, denoted by the pink triangle, occurs in a counterclockwise direction (pink dotted arrow), and regeneration does not occur at this end, indicating the lack of subunit incorporation at this end. (b) Schematic model of upper strand regeneration. The free subunit binds only to the plus end (cyan triangle) of the upper strand where the BB loop is exposed and does not bind to the minus end (pink triangle) where the EE surface is exposed. The convex surface represents the exposed BB loop, whereas the concave surface represents the EE surface. Notably, in the free subunit, the BB loop folds back onto the main body (Extended Data Fig. 3e,f,j,k). (c) Proposed free energy diagram illustrating the monomer incorporation into the upper strand shown in (b). Binding of the monomer to the minus end requires reorganization of the BB loop, which imposes a kinetic barrier.

In reference to our cryo-EM structure, regeneration occurs such that the exposed BB loop of the subunit at one end (designated the plus end) is bound by the EE surface of a free subunit. In contrast, the exposed EE surface of the subunit at the other end (minus end) is not bound by the BB loop of a free subunit (Fig. 5b). Structural comparison of TIR_MyD88_ in the rings with monomeric TIR_MyD88_ ^15^ revealed marked differences in the BB surface. Whereas in the monomers, the BB loop folds onto the body, in the rings, it projects to bind to the EE surface of a neighboring subunit (Supplementary Video 2 and Extended Data Fig. 3e,f). Thus, upon incorporation into a ring, the backbone of the BB loop undergoes a significant reorganization to settle into the intrastrand interface, and some of the side chains even translocate to the interstrand interface, thereby contributing to head-to-head dimerization. In contrast, the changes in the EE surface are subtle (backbone root mean square differences (RMSDs) of 7.9 Å and 0.82 Å for the BB and EE surfaces, respectively; Extended Data Fig. 3e). Based on this structural comparison, we speculate that the structural changes in the BB loop occur in concert with the interstrand interactions of TIR_MyD88_ prior to its binding to the EE surface. In this scenario, at the plus end of the upper strand (cyan triangle in Fig. 5), the BB loop is thought to be already rearranged in a form that facilitates its binding to the EE surface of another subunit, thus promoting subunit incorporation at that site. In contrast, the BB loop of a free subunits is not prepared for binding; thus, the EE surface at the minus end may be bound only rarely by free subunits (pink triangle in Fig. 5). Namely, the BB loop rearrangement is thought to be a kinetic barrier for the binding process (Fig. 5c). This difference in reactivity toward free subunits between the two open ends may be responsible for the observed counterclockwise elongation of the upper strand.

Because our observation is limited to the regeneration of the upper rings located on top of the preformed lower rings on the AFM stage, the exact process of double-stranded filament formation in either solution or in cells remains undetermined. However, the data indicate that the structural reorganization of the BB loop in free monomers is a slow process that may limit the establishment of intrastrand interactions. Thus, this step may impose a kinetic barrier to the initiation of TIR_MyD88_ self-assembly, preventing unwanted self-assembly and signaling unless the receptor is activated.

### HS-AFM reveals direct binding between the receptor TIR and the TIR_MyD88_ ring

Interactions between IL-1Rs/TLRs and MyD88 are mediated by homotypic interactions of their TIR domains. Since previous reports have demonstrated direct binding between GST-TIR_TLR2_ and TIR_MyD88_ ^18,32^, we sought to visualize this TIR-TIR interaction using HS-AFM. First, TIR_MyD88_ rings were prepared on the AFM stage (Fig. 6a), and GST-TIR_TLR2_ was added. As shown in Fig. 6b-d and Supplementary Video 3, topping of the particles on the rings was observed. We determined the frequency of such topping events in the HS-AFM movies recorded in the presence and absence of GST-TIR_TLR2_ as well as following the addition of either GST or TIR_TLR2_ alone (Fig. 6e). The findings demonstrated that these topping events were observed almost exclusively in the presence of GST-TIR_TLR2_. Because GST exists as a dimer, it is assumed that TIR_TLR2_ is forced to dimerize in the corresponding fusion proteins. Thus, these results indicate that only dimeric TIR_TLR2_ binds to TIR_MyD88_ rings.

**Fig. 6:**
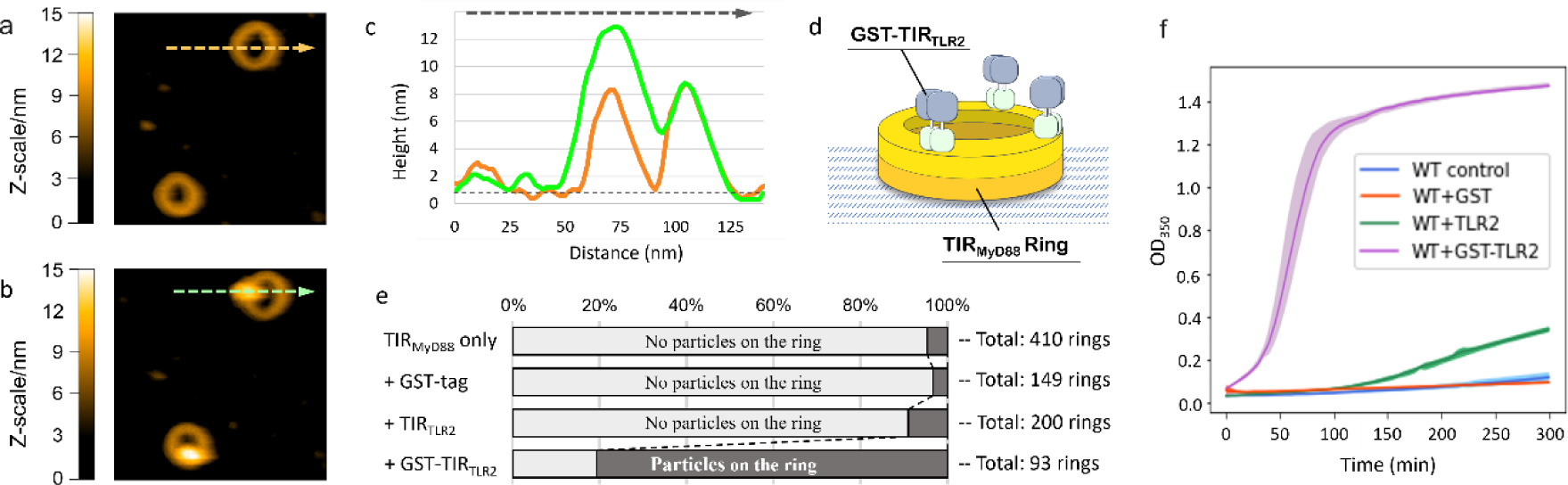
Direct visualization of GST-TIR_TLR2_ binding to TIR_MyD88_ rings. (a) Representative AFM image of preformed TIR_MyD88_ rings. (b) Rings bound by GST-TIR_TLR2_. Scanning area and scan speed in (a) and (b): 200 × 200 nm^2^ with 120 × 120 pixels and 500 ms per image. (c) Measurements of the cross sections represented by the dashed arrows in the AFM images in (a) (orange) and (b) (green). (d) Schematic model of the TIR_MyD88_ ring bound by GST-TIR_TLR2_. (e) Frequency of topping events. The dark areas of the bars indicate the percentages of rings with particles among the total number of rings. Rings with particles were frequently observed when GST-TIR_TLR2_ was added, whereas they were much less frequent when either GST or TIR_TLR2_ alone was added. (f) Turbidity assay of TIR_MyD88_ in the presence of GST-TIR_TLR2_, GST, or TIR_TLR2_. The addition of dimeric GST-TIR_TLR2_ markedly accelerated the increase in turbidity, but to a much lesser extent the addition of monomeric TIR_TLR2_.

While TIR_TLR2_ has been reported to form stable homodimers ^33,34^ and TLR2 can function as a homodimer under certain conditions ^35,36^, TLR2 is thought to function mainly as a heterodimer with TLR1 or TLR6 *in vivo*. Thus, to better reflect physiological conditions, we further tested another receptor TIR (TIR_TLR5_), which functions exclusively as a homodimer. GST-TIR_TLR5_ bound to preformed rings via the same mode as GST-TIR_TLR2_ (Extended Data Fig. 7), suggesting that the binding mode of these receptor TIRs is common. Importantly, both GST-TIR_TLR2_ and GST-TIR_TLR5_ bound to the upper surface of the rings, indicating that the interfilament surface of TLR_MyD88_ (Fig. 4a) functions as the binding site for the receptor TIRs.

### The self-assembly of TIR_MyD88_ is markedly accelerated by dimeric receptor TIRs

To determine the effect of GST-TIR_TLR2_ on the self-assembly of TIR_MyD88_ *in vitro*, we conducted a turbidity assay. When the concentration of TIR_MyD88_ was about 190 μM, the turbidity of the solution increased slowly after a lag time (blue line in Fig. 6f). However, a significant change was observed in the presence of a small amount of GST-TIR_TLR2_ (5% of the amount of TIR_MyD88_), with a rapid increase in turbidity observed (purple line in Fig. 6f). Addition of the GST tag alone did not result in this increase in turbidity, demonstrating that the accelerated self-assembly is caused by TIR_TLR2_. Interestingly, only marginal acceleration was observed when untagged TIR_TLR2_ (i.e., without the GST tag) was added (green line in Fig. 6f). TIR_TLR2_ has been reported to form both homodimers and tetramers at high concentrations ^33,34^. Given that GST is dimeric, TIR_TLR2_ is likely dimerized in its GST fusion protein. Thus, the data suggest that the dimerization (or at least the high local concentration) of TIR_TLR2_ plays an important role in the accelerated self-assembly of TIR_MyD88_. In addition, the amount of GST-TIR_TLR2_ required for the promotion of self-assembly was only 5% that of TIR_MyD88_, strongly suggesting that GST-TIR_TLR2_ is required only at the initial stage of assembly. Most likely, GST-TIR_TLR2_ assembles multiple TIR_MyD88_ subunits at one location to promote the nucleation process. After nucleation, TIR_MyD88_ self-assembly can occur rapidly, as previously hypothesized ^10,13^. Thus, these data demonstrate that dimeric receptor TIRs initiate the self-assembly of TIR_MyD88_ by promoting nucleation and that the effect of the dimers is substantially greater than that of the monomers.

## Discussion

### Tetramerization of TIR_MyD88_ is a plausible key step for its self-assembly

As mentioned above, the arrangement of four neighboring TIR_MyD88_ subunits in a parallelogram is thought to be geometrically suitable for promoting the oligomerization of its N-terminal DD_MyD88_ (Fig. 3b-d), because all the N-termini of TIR_MyD88_ protrude to one side in this arrangement (outer surface of the rings, Fig. 7a-c and Extended Data Fig. 3a). In DD_MyD88_ tetramers, four subunits are also arranged in a quasi-square configuration, and their C-termini protrude to one side (Extended Data Fig. 4). Therefore, by facing the N-terminal side of TIR_MyD88_ toward the C-terminal side of DD_MyD88_, a complete arrangement of the full-length MyD88 tetramer can be envisaged (Fig. 7c). Because of the intermediate domain ^4,8^, the relative position between TIR_MyD88_ and DD_MyD88_ is not fixed. However, on average, tetramerization of TIR_MyD88_ brings four N-terminal DD_MyD88_ subunits into proximity with the favorable orientation that leads to tetramerization of DD_MyD88_. Thus, the self-assembly of TIR_MyD88_ may promote the self-assembly of DD_MyD88_, which in turn initiates the recruitment of IRAK family members via DD-DD interactions. In this way, external signals can be transduced from receptors to IRAKs via Myddosome formation. MyD88 plays a central role in this process through its two distinct self-assembling domains, TIR_MyD88_ and DD_MyD88_. This signaling feature is reminiscent of a mechanism called signaling by cooperative assembly formation (SCAF), which is frequently found in innate immune and cell death signaling pathways ^37^.

**Fig. 7:**
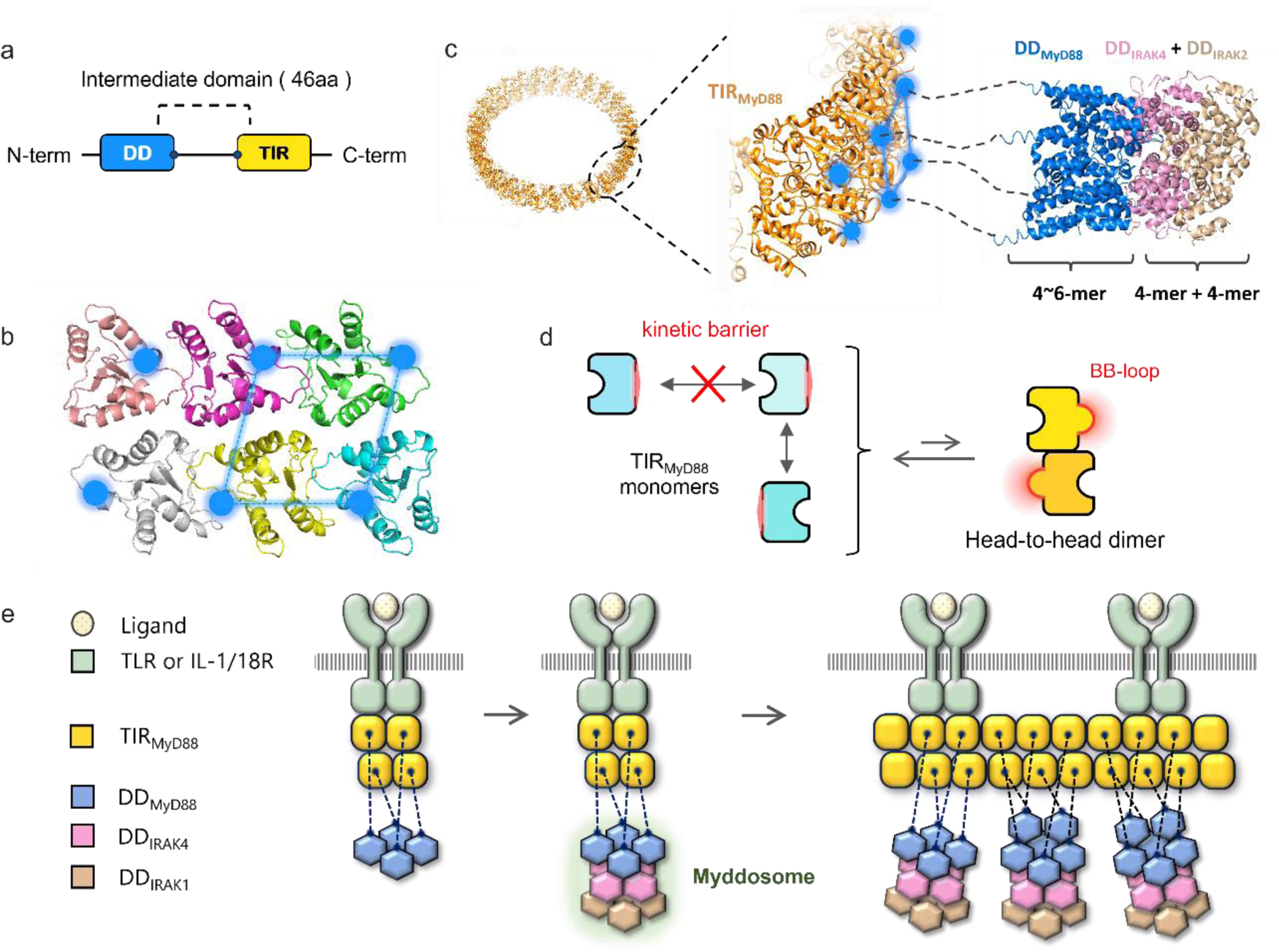
Schematic model of receptor-triggered oligomerization of MyD88. (a and b) Domain structure of MyD88 (a) and subunit arrangement in filaments (b). The filled cyan circles indicate the positions of the N-termini of TIR_MyD88_, to which DD_MyD88_ subunits are connected via linker residues. A parallelogram connecting the N-termini is shown in cyan. (c) Putative spatial relationship between the ring and the DD oligomer, in which TIR_MyD88_ (orange) and DD_MyD88_ (blue) are connected by the intermediate domain (dotted lines). DD_IRAK4_ (pink) and DD_IRAK2_ (light brown) are also shown. (d) Structural reorganization of the BB loop imposes a kinetic barrier for the intrastrand interactions, whereas head-to-head dimerization of TIR_MyD88_ induces rearrangement of the BB loop (red). (e) Schematic of receptor-triggered Myddosome formation and its clustering.

The tetrameric unit of TIR_MyD88_ described above is stabilized by inter- and intrastrand interactions, which are anticipated to work cooperatively and impart much greater stability than the trimeric or dimeric units. On the other hand, the establishment of intrastrand interactions requires a drastic rearrangement of the BB loop and its surrounding residues (Extended Data Fig. 3 and Supplementary Video 2). Given the polarity in strand elongation observed via AFM (Fig. 5), this rearrangement may impose a kinetic barrier that hampers the intrastrand interactions between free monomers. At physiological concentrations, this barrier is expected to prevent uncontrolled and unwanted intracellular self-assembly of TIR_MyD88_ (Fig. 7d). However, after the formation of a stable tetramer due to interactions with dimerized receptors, the incorporation of free subunits into the tetramer is expected to be markedly promoted because the tetramer can act as a stable platform containing BB loops suitable for binding, leading to rapid elongation of the antiparallel double-stranded filament.

As mentioned above, the rearrangement step of the BB loop into the binding form likely imposes a kinetic barrier for free monomers. On the other hand, the interstrand interface needs less structural reorganization, implying that head-to-head dimers form more readily, although their lifetime may be shorter (Fig. 7d). This transient dimerization can induce the BB loop rearrangement. Hence, in cells, it is possible that these dimers associate, through interactions with dimerized receptor TIRs, to form tetramers. In addition, TIR_MyD88_ subunits could bind as head-to-head dimers to the core tetramers (and preformed filaments) during further filament elongation. Although these possibilities await further investigation, intriguingly, the locations of oncogenic mutations that aberrantly facilitate MyD88 oligomerization supports this idea. Most of these mutations are in the BB loop and the head-to-head dimer interface (the interstrand interface, Supplementary Video 2 and Extended Data Fig. 3j). If a mutation lowers the kinetic barrier and stabilizes the head-to-head dimer of TIR_MyD88_, core tetramers and double-stranded filaments can readily form even in the absence of dimerized receptors, leading to constitutive and pathogenic signaling.

Based on the above, we propose that TIR_MyD88_ tetramerization functions as a checkpoint in signal transduction. AFM measurements revealed that both GST-TIR_TLR2_ and GST-TIR_TLR5_ specifically bind the upper surface of the rings (Fig. 6 and Extended Data Fig. 7). This spatial arrangement suggests that TIR_MyD88_ tetramers are formed below the dimeric receptor TIR (nucleation), from which filament elongation is initiated. It was previously demonstrated that upon stimulation of IL-1Rs in EL4.NOB cells, GFP-fused MyD88 assembled at receptor sites and formed clusters. The number of GFP-MyD88 molecules in the clusters was determined based on fluorescence intensity measurements, and IRAK4 was found to be recruited to MyD88 clusters containing an average of eleven MyD88 monomers. In contrast, smaller MyD88 clusters, containing an average of 3.2× MyD88 monomers, did not colocalize with IRAK4 ^13^. This finding suggested that the clustering of at least four MyD88 monomers is necessary for the recruitment of IRAK4. In addition, small oligomers (2-3 MyD88 monomers) were reported to be unstable, with a short lifetime, whereas larger oligomers had a longer lifetime and were more likely to coassemble with IRAK4. Thus, the authors proposed that the oligomer size of MyD88 is a “physical threshold” for induction of signaling. This hypothesis is fully consistent with the insight gained from our findings, which indicates that TIR_MyD88_ tetramerization is a critical step in the elongation of functional filaments (Fig. 7c). Considering previous data collectively with the findings of the current study, we propose a model for the functional self-assembly of MyD88 triggered by IL-1Rs and a subset of TLRs that do not require Mal (Fig. 7c-e). In this model, TIR_MyD88_ is first released from the autoinhibited state ^5^ and transiently forms symmetric head-to-head dimers via interstrand interactions (Fig. 7d). After membrane receptors dimerize, their TIR domains directly interact to assemble TIR_MyD88_, which promotes TIR_MyD88_ tetramerization (nucleation step, Fig. 7e, left). The stable tetramer functions as a platform for subunit incorporation, triggering filament elongation. This in turn promotes the oligomerization of DD_MyD88_ along the filament, ultimately leading to Myddosome formation at multiple sites on the filament (Fig. 7e). With this model, the formation of multi-Myddosome clusters is also feasible ^38^.

### Functional form of the double-stranded filaments of TIR_MyD88_

Because the cylindrical fibers of TIR_MyD88_ take much longer to form than the rings (Fig. 1a) and the interfilament mutants do not significantly affect the dominant negative inhibitory effect (Fig. 4b), they may not represent the functional state in the cell. In addition, the average number of GFP-MyD88 molecules that accumulated at IL-1Rs recruiting IRAK4 was previously found to be 11 on average ^13^, whereas the number of subunits in a double-layered ring of TIR_MyD88_ was at least 52. Therefore, relatively short free double-stranded filaments are thought to constitute the functional oligomeric state of TIR_MyD88_. The reported stoichiometry of DD_MyD88_ and DD_IRAK4_ in the complexes varies, with reported ratios of 6:4, 7:4 and 8:4 ^39,40^. Our double-stranded filament of TIR_MyD88_ is compatible with any of these stoichiometries since DD_MyD88_ subunits, which are connected to TIR_MyD88_ subunits outside of the parallelogram formed by four TIR_MyD88_ subunits, can also participate in DD-DD oligomerization (Fig. 7c). The incorporation of two, three, or four additional DD_MyD88_ subunits, which correspond to the aforementioned stoichiometries, further stabilizes the DD_MyD88_ oligomer, resulting in a more robust platform for IRAK4 recruitment.

### An evolutionarily conserved self-assembly mode of the TIR domain

Recently, self-assembled structures of the TIR domains of plant and bacterial proteins have been reported ^41–44^, in which BB loops commonly play a crucial role in the formation of higher-order structures ^45^. Among these structures, the filament of TIR-STING, an antiphage effector protein of *Sphingobacterium faecium*, is noteworthy ^41^, as the subunit arrangement in this filament is approximately the same as that in the double-stranded filament of TIR_MyD88_, despite the evolutionarily distant relationship (Extended Data Fig. 8a,b). Homodimeric TIR-STING forms filaments upon binding of the nucleotide second messenger c-di-GMP at the STING domain. TIR_STING_ subsequently becomes enzymatically active to cleave NAD^+^. Impressively, to acquire enzymatic activity, TIR_STING_ needs to form at least one tetramer, in which the active site is created through the interaction between the BB loop of one subunit and the EE surface of its neighboring subunit. The formation of TIR tetramers has also been reported to trigger the effector functions of other TIR-containing proteins. hSARM1 ^43^, ROQ1 (a protein related to plant immunity) ^42^, and RPP1 ^44^ are examples (Extended Data Fig. 8c-h), although their subunit arrangements differ from that of TIR_MyD88_ and TIR_STING_ assemblies. Thus, tetramerization is most likely a conserved feature required for the function of TIR-containing proteins. Another noteworthy feature of the TIR domains of hSARM1, ROQ1, and RPP1 is that they commonly utilize a surface that we defined as the interfilament surface for tetramerization (Extended Data Fig. 8c,d,e), indicating the functional relevance of this surface. Consistent with this idea, our AFM measurements identified the interfilament surface as the binding site for dimeric receptor TIRs (Fig. 6 and Extended Data Fig. 7). Hence, it may be valuable to focus on the interfilament surface to determine the mode of interaction between the receptor and MyD88. The functional importance of TIR tetramerization demonstrated in these proteins, together with the dramatic increase in thermostability due to cooperative interactions in tetramerization, supports our model in the assertion that TIR_MyD88_ tetramerization is the critical step (nucleation) for the elongation of double-stranded filaments, which is controlled by receptor dimerization (Fig. 7).

In summary, we determined the structure of TIR_MyD88_ filaments using cryo-EM. The architecture provides a geometric explanation for the regulated recruitment of downstream IRAKs via antiparallel double-stranded filament formation as well as insights into disease mutations. Our HS-AFM and turbidity assay results greatly advanced our mechanistic understanding of the receptor-triggered oligomerization of MyD88. The structure and molecular scheme proposed herein will prompt further mechanistic analysis of this key adaptor protein and increase our understanding of Myddosome function.

## ACKNOWLEDGMENTS

We thank Naotaka Tsutsumi (Okayama University), Takayuki Kato (Osaka University) and Yudai Ito (Kyoto University) for their scientific advice or technical help. We also thank Toshio Ando (Kanazawa University) for the technical support of HS-AFM and Keiko Okamoto-Furuta and Haruyasu Kohda (Kyoto University) for technical assistance in electron microscopy. AFM data collection was supported by the World Premier International Research Center Initiative of the Ministry of Education, Culture, Sports, Science and Technology, Japan. This work was supported by CREST, Japan Science and Technology (JST) Agency (JPMJCR1762) and JSPS KAKENHI (23H02421) to H.T., JST SPRING (JPMJSP2110) to K. K., Health and Labour Science Research Grants for Research on Intractable Diseases from the Ministry of Health, Labour and Welfare (23809849 and 23809798) and Grants of the Japan Agency for Medical Research and Development (AMED) (22582994, 23808661 and 24015303) to H.O. This work was partly supported by Bio-SPM Collaborative Research (Kanazawa University) and Research Support Project for Life Science and Drug Discovery (Basis for Supporting Innovative Drug Discovery and Life Science Research (BINDS)) from AMED under Grant Number JP22ama121003.

## Author contribution

Conceptualization, K.K., M.U. and H.T.; data curation, K.K., K.I. and N.S.; formal analysis, K.K., K.I. A.N. and H.T.; funding acquisition, H.O. and H.T.; investigation, K.K., M.U., T.M., F.M., R.Y., Y.T. and H.K.; methodology, K.K., H.O. and H.T.; project administration, H.T.; resources, K.N., H.O., H.K. and H.T.; software, F.M. and N.K.; supervision, H.T.; validation, K.K. and H.T.; writing—original draft preparation, K.K. and H.T.; writing—review and editing, K.K., K.I., N.K., K.N., H.O., A.N., H.K. and H.T

---

**Extended Data Fig. 1:**
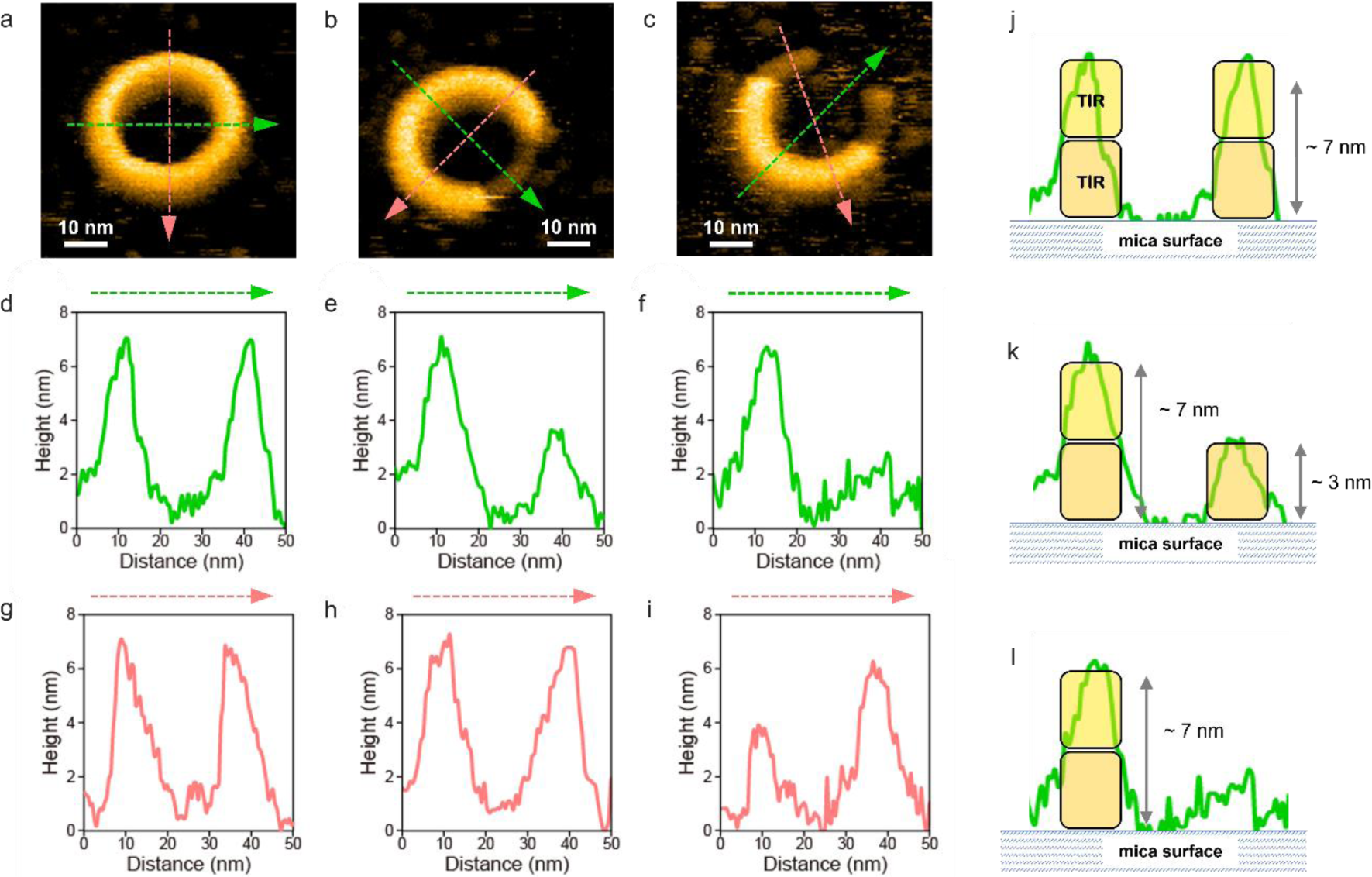
The TIR_MyD88_ ring is double layered. Related to Fig. 2. (a-c) Representative AFM images of the rings in the disintegration process. Each image was extracted at 13.84 s (a), 17.24 s (b) and 49.40 s (c) in Supplementary Video 1. Scan area: 60 × 60 nm^2^ with 120 × 120 pixels, Z-scale, 8.4 nm. Scan speed: 600 ms per image. (d-i) Sectional views of the green and pink arrows in the AFM images in (a), (b) and (c), respectively. The diameter of the rings was approximately 29 nm, and the height of the double layered rings was approximately 6.5 nm. In the partially lacking ring, the heights of lacking regions were approximately 3.2 nm (e,i) or less than 2 nm (f). (j-l) Schematic representations of subunit arrangements in (d), (e), and (f), respectively.

**Extended Data Fig. 2:**
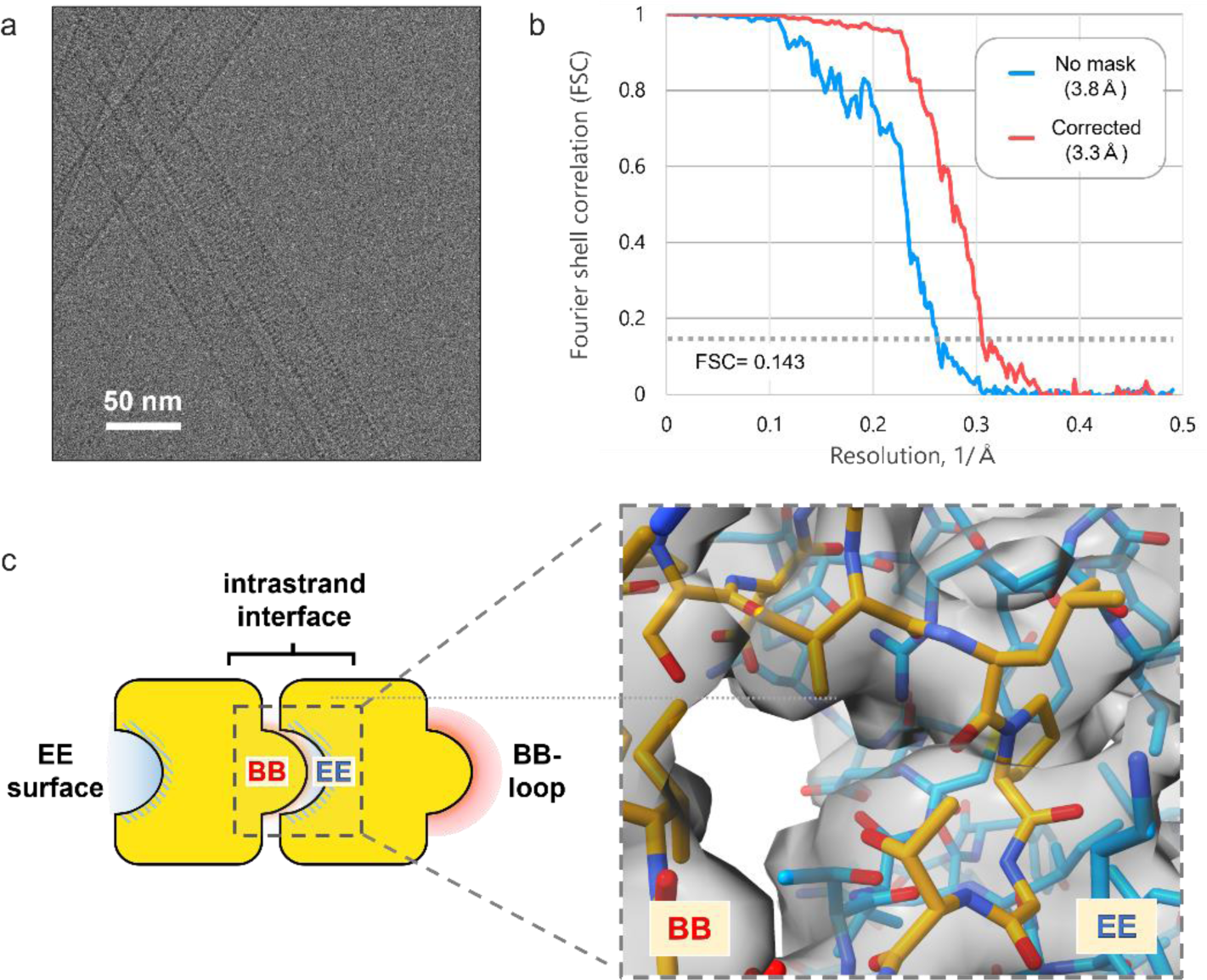
Cryo-EM imaging and the evaluation of the density map. Related to Fig. 3. (a) A typical motion-corrected cryo-EM micrograph of the cylindrical fibers formed by stacked rings. (b) Fourier shell correlations (FSCs) for evaluating the resolution calculated by CryoSPARK v3.2.0 (Punjani et al. 2017). FSCs between two 3D structures from each half of the dataset with or without masking are presented. Based on the golden standard criteria, the resolution is 3.3 Å with a threshold of 0.143. (c) A cryo-EM map around the intrastrand interface. The view is from the outside of the cylindrical fiber, focusing on the BB loop (orange stick) and EE surface (blue stick), which consists of βD-strand, βE-strand, and αE-helix.

**Extended Data Fig. 3:**
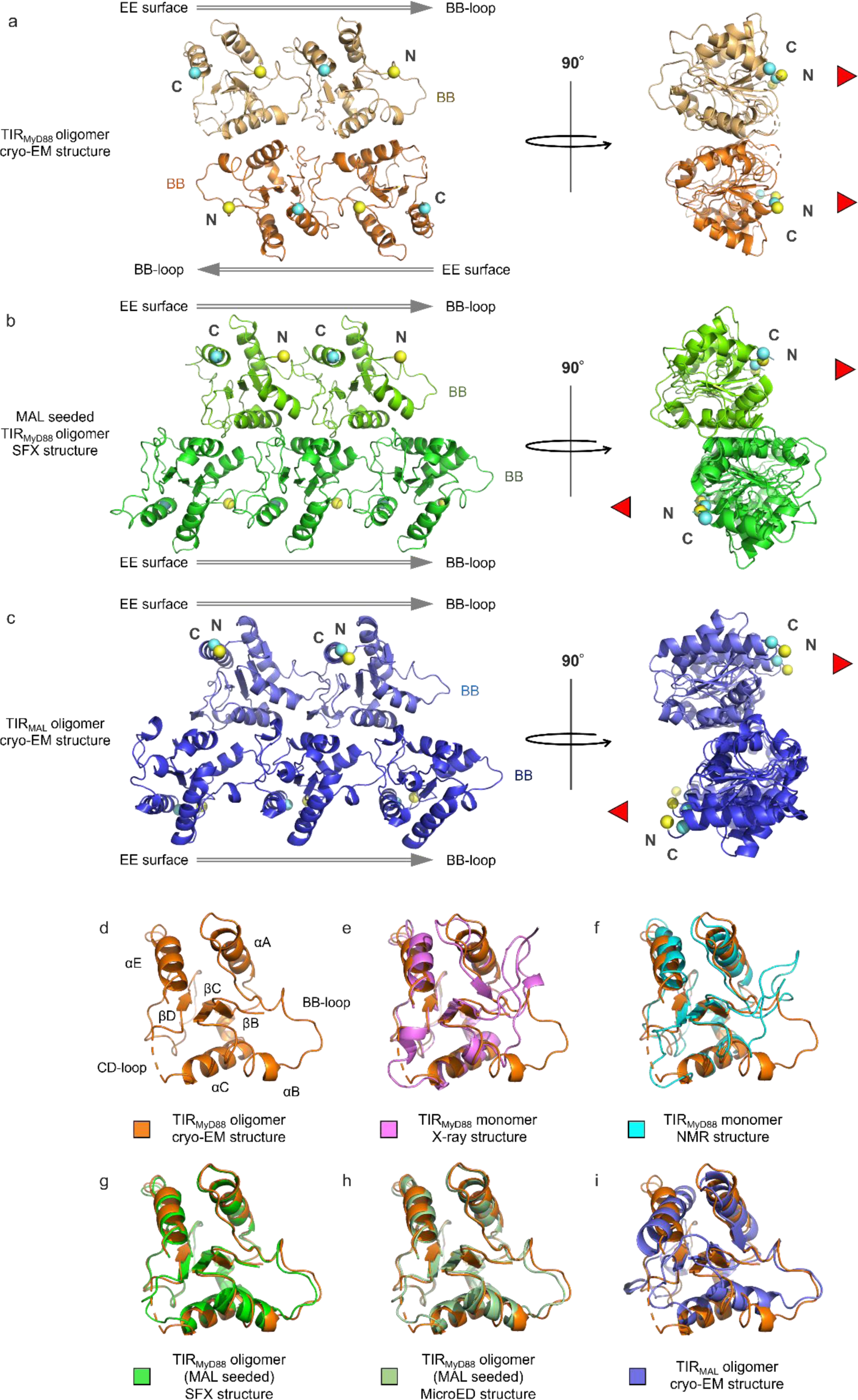
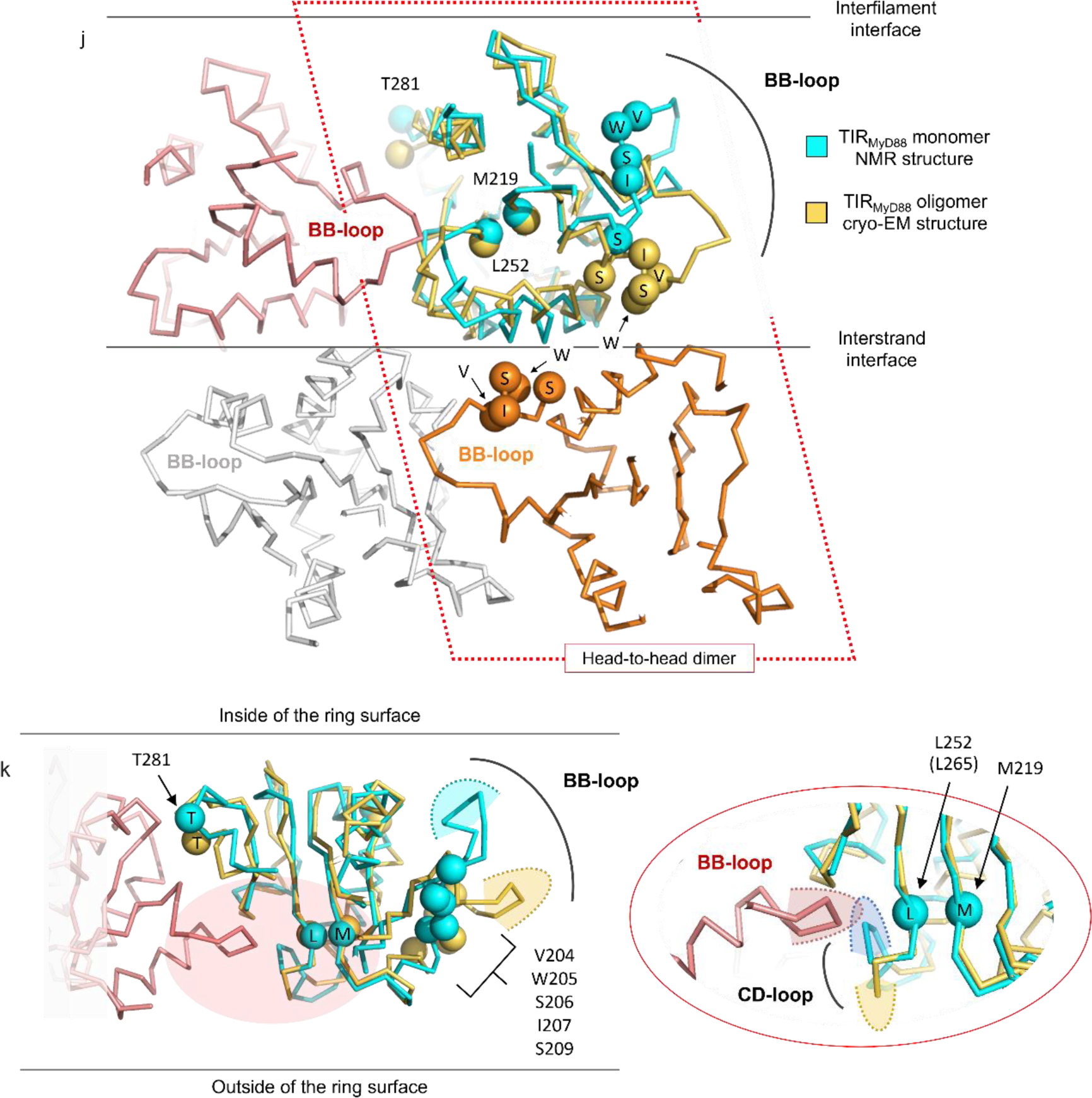
Structural comparison of TIR_MyD88_ in various states. Related to Fig. 3. (a-c) The cryo-EM structure in this work (a) and Mal-induced crystal structure (b) (Clabbers et al. 2021) of the TIR_MyD88_ oligomer are of a double-stranded linear assembly. In both cases, each strand consists of tandemly arrayed subunits that associate via the intrastrand interactions. However, the alignment of the two strands is completely opposite. In the Mal-induced oligomer, two TIR_MyD88_ strands are joined in a *parallel* orientation (b) identical to that of the Mal fibril (c) (Ve et al. 2017). In contrast, in the cryo-EM structure, the two strands are in an *antiparallel* orientation (a). Additionally, in the Mal-induced oligomers, each subunit in one of the strands is wedged between two adjacent subunits in the other strand, whereas in the cryo-EM structure, one subunit in one strand is in complementary and symmetrical contact with one subunit in the other strand. As a result, in (a), all the N-termini, where DD_MyD88_ connects, are located on one side as indicated by the red triangles. In contrast, in (b) and (c), the N-termini project into two opposite sides of the filament as indicated by the red triangles. The N- and C-termini are depicted in yellow and cyan spheres, respectively. (d-i) Structural comparison of TIR_MyD88_ determined in this study (in orange) with structures from other studies. The structure of the BB loop is significantly different from the monomeric states ((e) PDB ID: 4EO7 in pink and (f) 2Z5V in cyan) (Ohnishi et al. 2009; Snyder et al. 2013), though similar to other oligomeric states of TIR_MyD88_ ((g) PDB ID: 7BER in green and (h) 7BEQ in pale green) and TIR_Mal_ ((i) PDB ID: 5UZB (purple)) (Ve et al. 2017; Clabbers et al. 2021). In (e), root mean square differences (RMSDs) for the BB and EE surfaces aligned based on the protein backbone were 7.9 Å and 0.82 Å, respectively. (j-k) Detailed comparison of the cryo-EM structure of oligomeric state of TIR_MyD88_ in the double-stranded filament with its monomeric state. See Supplementary Video 2 for a morphing movie showing the structural difference. Views from the outer surface of the double-layered ring (j) and the interfilament interface (k) are depicted. In (k), the region encircled by the red ellipse in the left figure is enlarged in the right. The black curved lines outline the occupied space of the tips of the BB loop and CD loop. The areas enclosed by the dotted curves represent the tips of the BB loop and CD loop in monomeric or oligomeric state. During self-assembly, the BB loop undergoes reorganization, leading to both the intra- and interstrand interactions. Gain-of-function mutation residues (referred to as V204, W205, S206, I207, S209, M219, L252, and T281 in our study) associated with aggressive B-cell lymphoma are represented as spheres labeled with single-letter amino acid codes (Ngo et al. 2011; Yu et al. 2018; de Groen et al. 2019). In the antiparallel double-stranded filament, residues within the BB loop (V204-I207 and S209) are situated at the interstrand interface, suggesting their role in head-to-head dimerization. Three other residues (M219, L252, and T281) are positioned at or close to the intrastrand interface. Thus, all of the oncogenic mutations mentioned above occur in positions crucial for maintaining the antiparallel double-stranded filament structure. This suggests that these oncogenic mutations strongly promote filament formation (O’Carroll et al. 2018) by enhancing either inter- or intrastrand interactions, or both, resulting in unregulated downstream activation.

**Extended Data Fig. 4:**
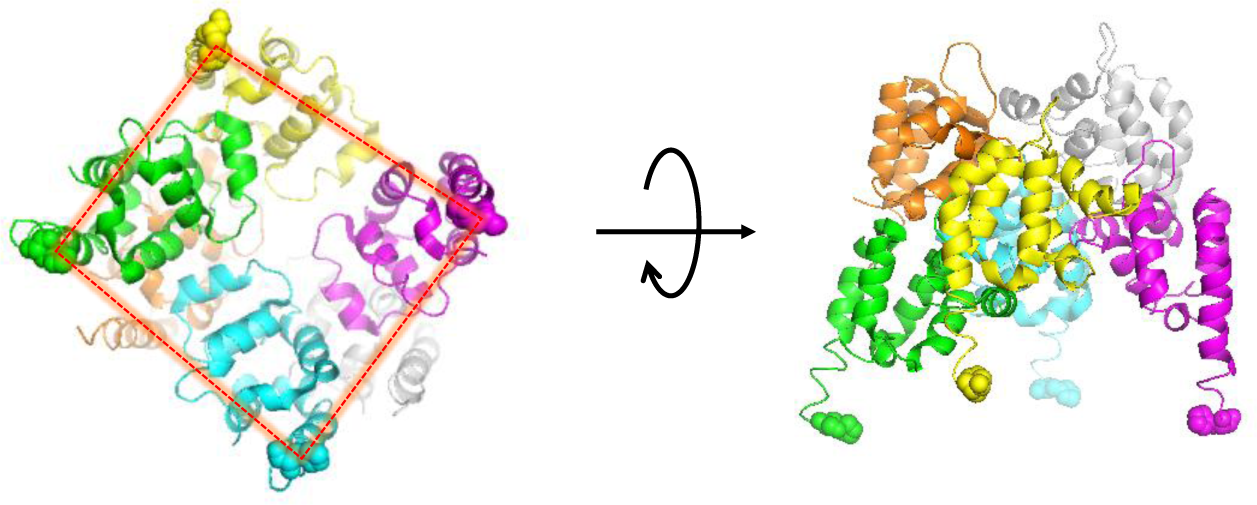
Four DD_MyD88_ subunits make a quasi-square in the helical oligomer. Related to Fig. 3 and 7. Six DD_MyD88_ subunits extracted from the helical complex with DD_IRAK4_ and DD_IRAK2_ are shown (PDB ID: 3MOP) (Lin, Lo, and Wu 2010). The sphere models indicate the C-terminal residues of four DD_MyD88_ subunits in the bottom layer, to which TIR_MyD88_ are connected via linker residues (the intermediate domain). A quasi-square connecting the C-termini of DD_MyD88_ is represented by a red dotted line.

**Extended Data Fig. 5:**
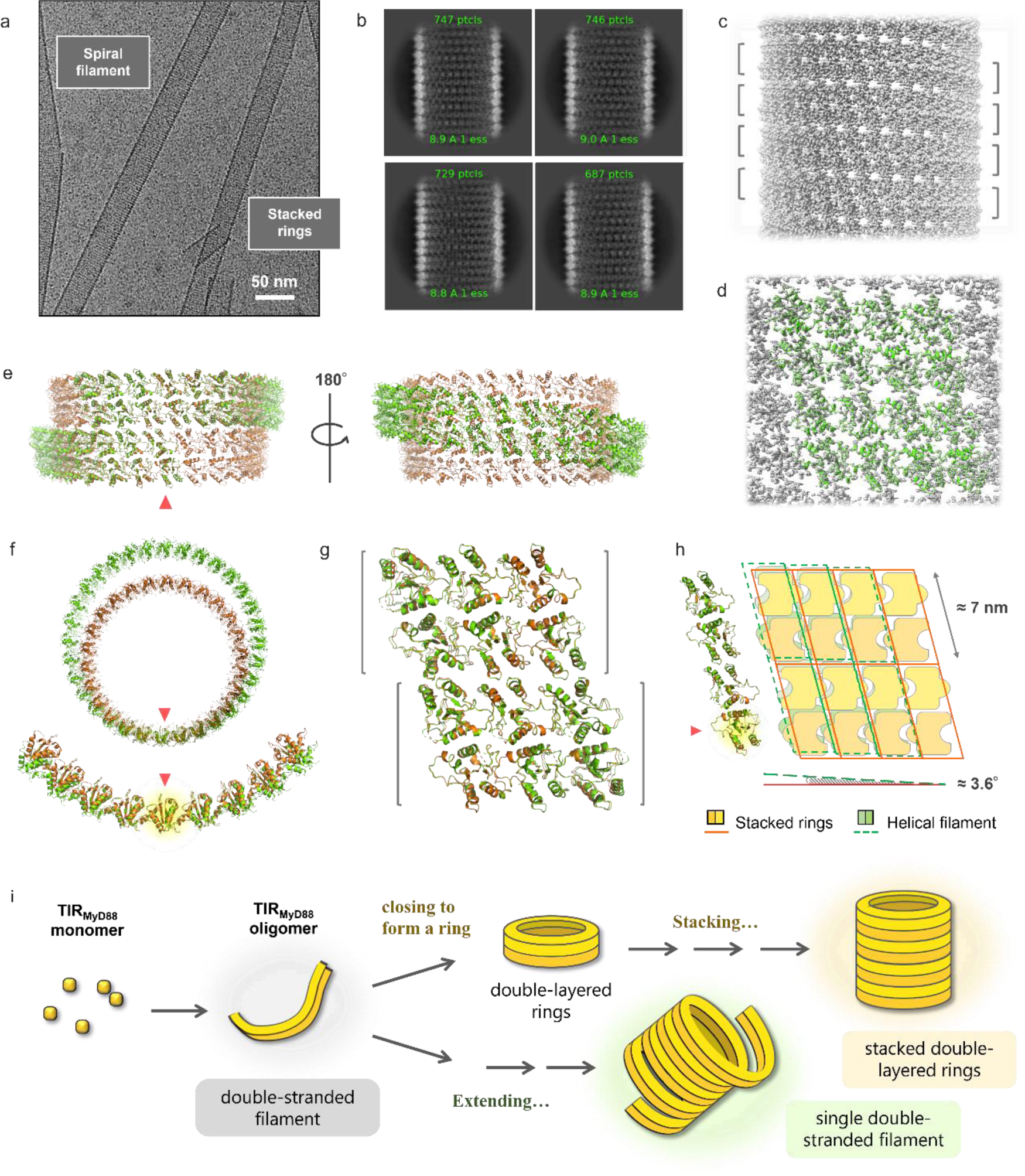
Spiral cylinder; overall and subunit arrangement. Related to Fig. 3. (a) An example of a cryo-EM micrograph showing two distinct cylindrical fibers of different diameters; stacked rings (see Fig. 3 and Extended Data Fig. 2) and spiral filament. (b) Representative 2D class averaged images of the spiral filament. (c) Cryo-EM density map of the cylindrical fiber formed by the spiral double-stranded filament. Gray brackets indicate the width of the double-stranded filament. (d) Molecular models fitted to the EM map. (e, f) A comparison of subunit arrangements in the stacked-ring filament (orange) and the spiral filament (green) is depicted. Side (e) and top views (f) of the two cylindrical fibers are presented. The two cylinders are aligned using a subunit indicated by red triangles. (g) Superposition of 12 subunits in the stacked-ring and the spiral filament. Gray brackets indicate the width of the double-stranded filaments (i.e. double-layered ring). (h) Comparison of the subunit arrangements in the two filaments (side view). The subunit arrangement is essentially the same for both filaments. In particular, the vertical positioning of the subunits is nearly identical (left: four vertically arrayed subunits are aligned with the subunit indicated by the red triangle). However, their lateral positioning slightly differs, resulting in a tilt of approximately 3.6 degrees. This tilt creates the difference, either the stacked rings or the spiral filament. (i) Schematic of stacked rings and spiral cylinder formation. The spiral cylinder is formed from a single antiparallel double-stranded filament wound into a spiral. The two types of cylinders are formed from a common filament. The difference is whether the filament closes to form a ring or extends infinitely in a spiral.

**Extended Data Fig. 6:**
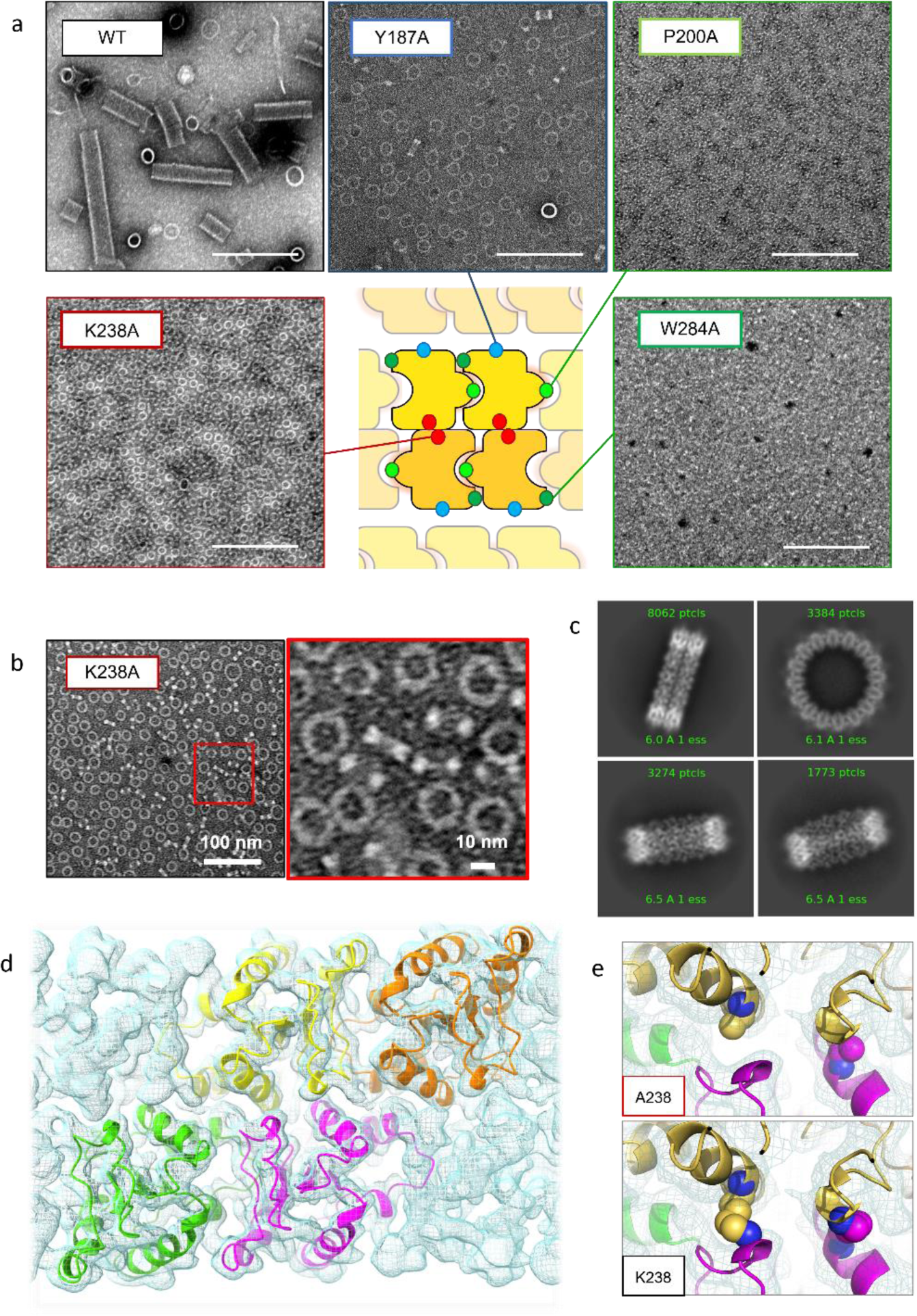
Self-assemble propensity of various TIR_MyD88_ mutants. Related to. **Fig. 4** (a) TEM images of WT TIR_MyD88_ or mutants of TIR_MyD88_. The scale bar in (a) represents 200 nm. The positions of the mutated residues are shown in the cartoon. (b) TEM and (c) class-averaged cryo-EM images of the K238A rings. (d) Four molecular models of K238A TIR_MyD88_ fitted into the K238A EM map (FSC resolution was 3.2Å) are shown. (e) A closer look at the sidechains (spheres) of two A238 residues from two subunits in (d) is shown (top). Replacing these Ala residues with Lys (as in the WT sequence) would result in collisions with the adjacent subunits (bottom). As shown in (a), mutants of TIR_MyD88_ with the P200A or W284A (intrastrand mutation) completely failed to self-assemble, while the Y187A mutant (interfilament mutation) formed rings similar to those formed by WT TIR_MyD88_, but largely failed to form cylindrical fibers. The K238A mutant (interstrand mutation) formed rings but not cylindrical fibers. Notably, the diameter of the K238A mutant rings is significantly smaller, ranging from 14-25 nm (b), compared to the WT rings (22-38 nm). In fact, cryo-EM analysis of the K238A mutant rings revealed that the subunit arrangement is different from the WT TIR_MyD88_ ring (d). Importantly, such an arrangement is incompatible with WT TIR_MyD88_ due to steric clash of the K238 sidechains with adjacent subunits as shown in (e). This means that ectopically expressed K238A TIR_MyD88_ molecules in 293T cells as in Fig. 4b are essentially unable to co-assemble with endogenous WT MyD88, even if the K238A TIR_MyD88_ molecules can form homooligomers on their own. Therefore, dominant negative inhibition by the K238A mutant is not expected, which is consistent with the result shown in Fig. 4b.

**Extended Data Fig. 7:**
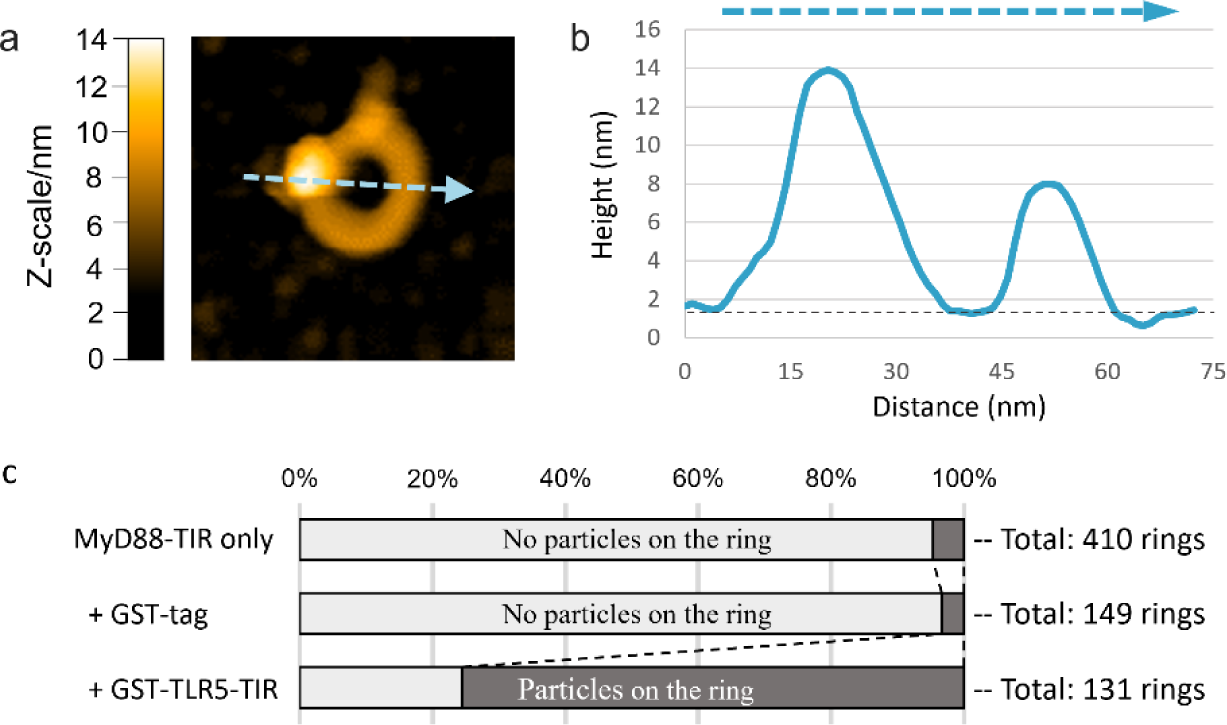
Direct visualization of the binding of GST-TIR_TLR5_ to the TIR_MyD88_ rings. Related to Fig. 6. (a) A representative averaged AFM image of WT rings bound by GST-TIR_TLR5_. Scan area: 100 × 100 nm^2^ with 100 × 100 pixels. Scan speed: 230 ms per image. (b) A sectional view along the dashed arrow in the AFM image in (a). (c) Frequency of topping events. The dark areas of the bars indicate the percentages of rings with particles among the total number of rings. Rings with particles were frequently observed when GST-TIR_TLR5_ was added, whereas they were not observed when only either GST or TIR_TLR5_ was added. While TIR_TLR2_ has been reported to form stable homodimers (Tao et al., 2002; Xu et al., 2000) and TLR2 has been proposed to function as a homodimer (Bagheri et al., 2014; Zheng et al., 2015), TLR2 is generally thought to function as heterodimers with TLR1 or TLR6 during signaling *in vivo*. Also, TLR1/2 and TLR2/6 employ the other adaptor protein, Mal, in addition to MyD88 (Horng et al., 2002). To gain more direct insight into biological relevance, we examined TIR_TLR5_ for MyD88 binding, because TLR5 functions exclusively as a homodimer and does not require Mal for signaling *in vivo* (Horng et al., 2002; Voogdt et al., 2018). The HS-AFM measurement showed that GST-TIR_TLR5_ also binds to preformed rings in the same manner as GST-TIR_TLR2_.

**Extended Data Fig. 8:**
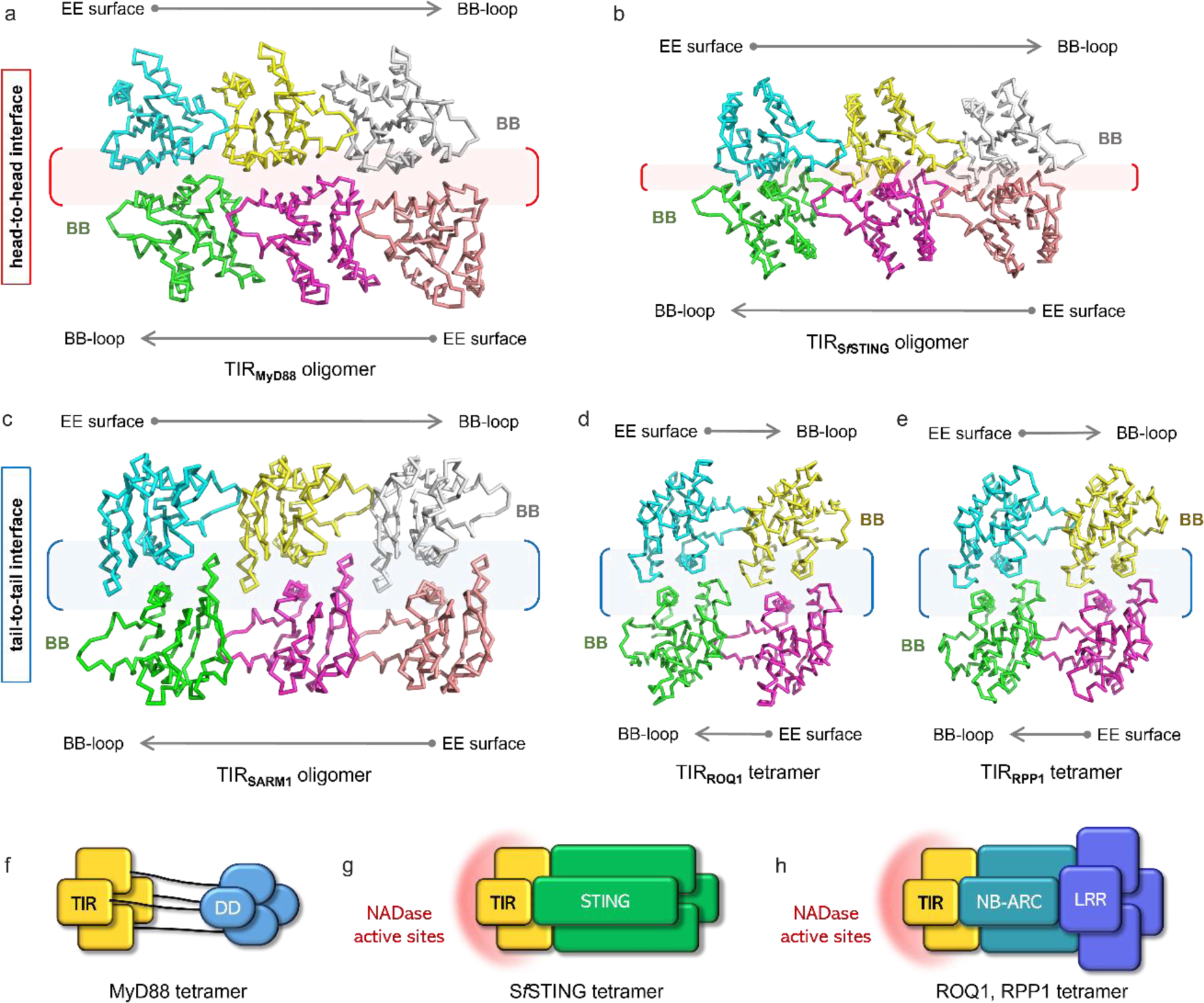
Comparison of the subunit arrangement in the double-stranded filament of TIR_MyD88_ with other TIR assemblies. (a-b) The subunit arrangement in the antiparallel double-stranded filament of TIR_MyD88_ determined in this work (a) and the *Sf* TIR-STING filament (b) from the antiphage effector protein of *Sphingobacterium faecium* are shown (Morehouse et al. 2022). The homo-dimeric TIR-STING forms filaments upon binding to the nucleotide second messenger c-di-GMP at the STING domain. TIR_STING_ subsequently exhibits enzymatic activity to cleave NAD^+^. In the filaments, homodimers of TIR_STING_ are arrayed laterally and the resulting subunit arrangement is topologically identical to that of TIR_MyD88_. The lateral interactions between the subunits are mediated through the BB loop and the EE surface, just like the intrastrand interactions in the TIR_MyD88_ filaments. In both TIR_MyD88_ and TIR_STING_ filaments, the two strands are antiparallel, interacting through the interstrand interface. However, in the TIR_STING_ filament, the interstrand interactions are much more extensive, occupying ∼16% (∼1250 Å²) of the total surface area of one subunit, compared to those in the TIR_MyD88_ filament, which occupy ∼5% (∼440 Å²). (c) The subunit arrangement in the TIR_hSARM1_ filament reveals that the two strands align antiparallel and associate through the interfilament interface (Shi et al. 2022). (d-e) TIR_ROQ1_ and TIR_RPP1_ both form tetramers rather than extended filamentous assemblies (Martin et al. 2020; Ma et al. 2020). Nevertheless, the arrangement of the tetrameric subunits is essentially the same as that of the TIR_hSARM1_ filament in (c). (f-h) Schematic representation of tetramers of TIR-containing proteins, MyD88 (f), *Sf* TIR-STING (g), and ROQ1/RPP1 (h). It has been reported that the tetramerization of TIR_STING_, TIR_ROQ1_, and TIR_RPP1_ occurs cooperatively with the tetramerization of their co-existing effector domains (g and h). Hence, the cooperative character of tetramerization is likely conserved in these evolutionarily distant TIR-containing proteins. This strongly supports the idea that tetramerization of TIR_MyD88_ (f), triggered by activated TLRs/IL-1R, promotes self-assembly of DD_MyD88_, which leads to the assembly of downstream IRAKs.

**Supplementary Video 1**

HS-AFM movie of disintegration and regeneration of TIR_MyD88_ rings. Related to Fig. 5. Scan area: 60 × 60 nm^2^ with 120 × 120 pixels. Scan speed: 600 ms per image.

**Supplementary Video 2**

Structural comparison of TIR_MyD88_ in monomeric and oligomeric states. Related to Fig. 7d, Extended Data Fig. 3j and 3k. The spheres represent Cα atoms of amino acid residues that are frequently mutated in aggressive B-cell lymphoma.

**Supplementary Video 3**

HS-AFM movie of GST-TIR_TLR2_ binding to TIR_MyD88_ rings. Related to Fig. 6. Scan area: 200 × 200 nm^2^ with 120 × 120 pixels. Scan speed: 500 ms per image.

**Supplementary Table 1:**
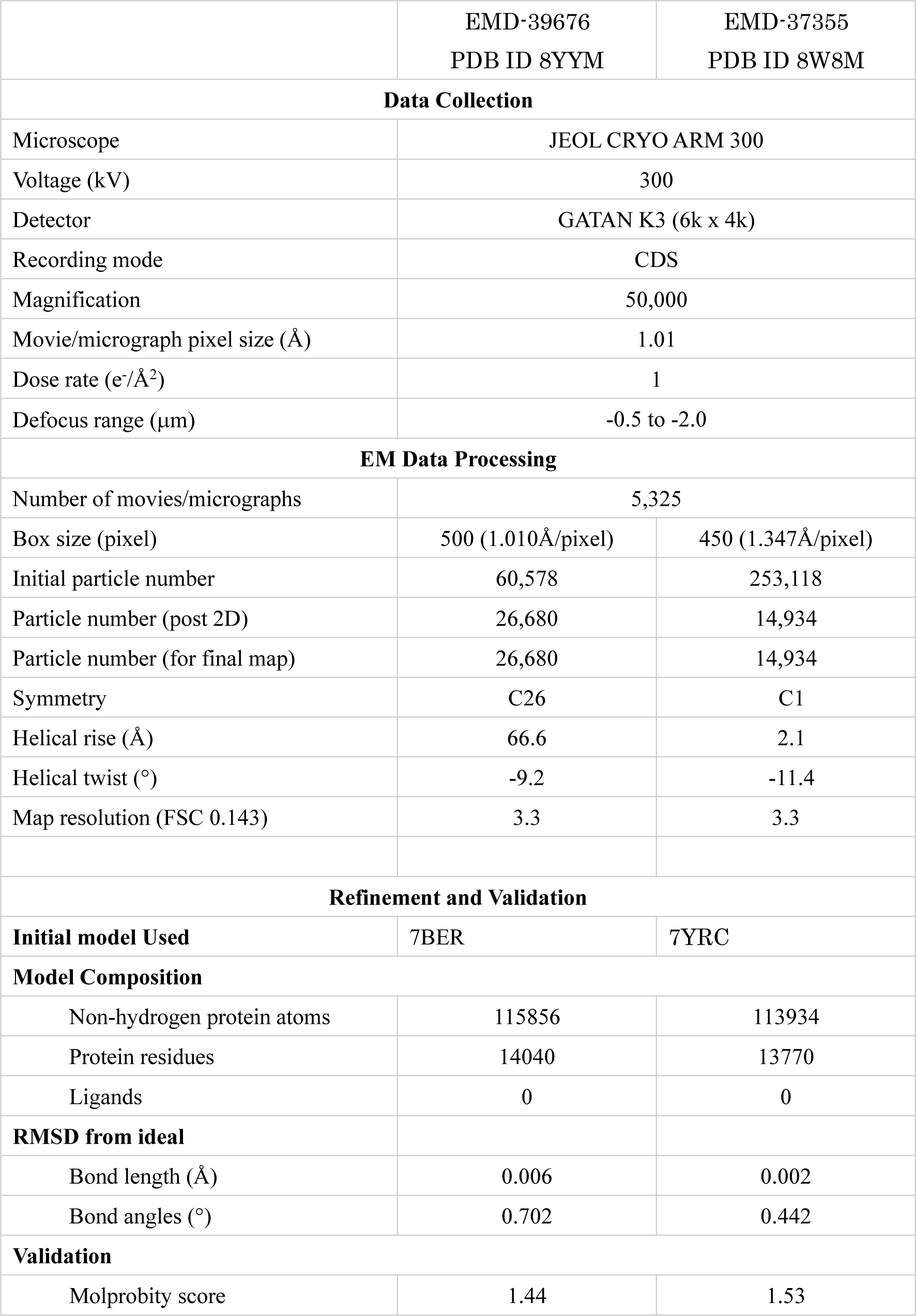
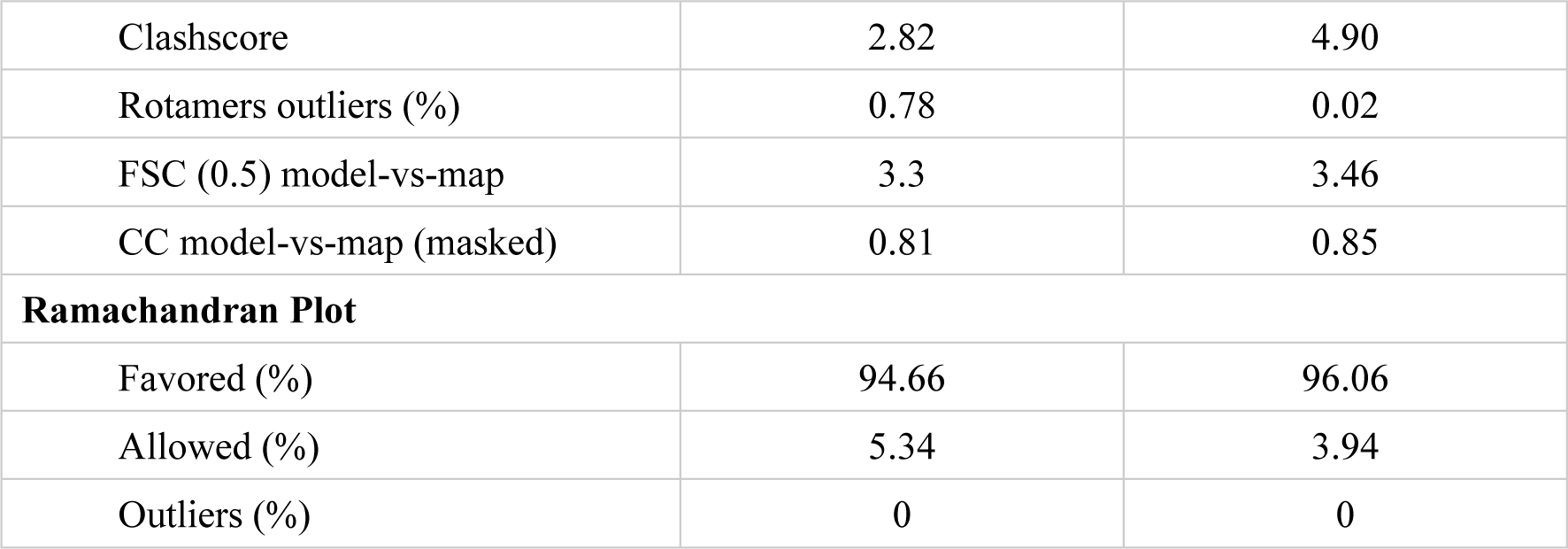
Cryo-EM data collection and structure refinement. Related to Fig. 3 and Extended Data Fig. 5.

## Materials and Methods

### Construction of expression vectors

The coding region of TIR_MyD88_ (MyD88, amino acid residues 153-296) was cloned into a pET-28a vector (Novagen), which was engineered with a 6×His-tagged streptococcal protein GB1 domain (GB1) for protein expression and purification. TIR_MyD88_ (residues 148-296) with an N-terminus myc-epitope was also cloned into pcDNA3.1+ vector (Invitrogen) for the luciferase activity assay. The constructs for the derivatives, such as the alanine-substituted derivatives, were prepared by site-directed mutagenesis. The TLR2 gene encoding the TIR domain (amino acid residues 633–784) and the TLR5 gene encoding the TIR domain (amino acid residues 685–858) were each cloned separately into a pGEX-6P-1 vector ^1^. IL-18Rβ construct tagged at the C-terminus with an AU1-epitope was also cloned into pcDNA3.1+. The NF-κB luciferase reporter vector (pGL4.32-luc2P/NF-kappaB-RE/Hygro) and the Renilla luciferase reporter vector (pGL4.74-hRluc/TK) were purchased from Promega.

### Protein expression and purification

For experimental samples of TIR_MyD88_, the expression vector for the His-GB1 tag fusion TIR_MyD88_ was transformed into *Escherichia coli* BL21(DE3) cells. The cells transformed with the His-GB1-tagged fusion TIR_MyD88_ vector were grown in LB medium. The His-GB1-tagged fusion TIR_MyD88_ was first purified with His-tag affinity column chromatography (cOmplete™ His-Tag Purification Resin: Roche). After removal of the His-GB1 tag with GST-HRV3C, the protein was further purified with cation exchange chromatography (HiTrap™ SP HP: Cytiva) and size-exclusion chromatography (HiLoad® 16/600 Superdex® 75 pg: Cytiva).

For experimental samples of GST-TIR_TLR2/TLR5_, the expression vectors were transformed into *Escherichia coli* BL-21 (DE3) (Novagen). The expressed GST fusion proteins were purified by Glutathione Sepharose™ 4 FF (Cytiva) affinity chromatography and size-exclusion chromatography (HiLoad® 16/600 Superdex® 75 pg: Cytiva) ^1^. For preparing TIR_TLR2_, before size-exclusion chromatography, the GST-tag was removed by digestion with GST-HRV3C followed by anion exchange chromatography (HiTrap™ Q HP: Cytiva).

### Dominant negative assay with NF-κB reporter gene activity

HEK293T cells were cultured in Dulbecco’s Modified Eagle Medium (high glucose-containing D-MEM, Invitrogen) supplemented with 10% heat-inactivated fetal bovine serum (SIGMA-ALDRICH), penicillin (100 U/mL) and streptomycin (100 µg/mL). All cells were incubated at 37°C in a humidified atmosphere of 5% CO_2_. For the reporter gene assays, HEK293T cells were transfected with 50 ng per well of pcDNA3.1+ mock vector or pcDNA3.1+ myc-TIR_MyD88_ (wild-type or mutants) in 96-well plates using Lipofectamine 2000 (Invitrogen) according to the manufacturer’s instructions. The NF-κB luciferase reporter, Renilla luciferase reporter vectors and pcDNA3.1+ IL-18Rβ were co-transfected. After transfection, cells were incubated for 24 hours, then stimulated with recombinant human IL-18 (20 ng/mL) for 6 hours. Luciferase reporter activity was analyzed using the Dual-Luciferase Reporter Assay System (Promega) as per the protocols.

### Turbidity assay

Self-assembly of TIR_MyD88_ was monitored at 37°C in 20 mM HEPES pH 7.0, 50 mM NaCl, 5 mM DTT by the change in turbidity (optical density at 350 nm) in a UV-Star® Microplate (Greiner Bio-One). The TIR_MyD88_ samples without or with TIR_TLR2/5_ were incubated by shaking at 108 rpm for 5 hours with a SPARK® multimode microplate reader (TECAN), attached with a Humidity cassette to keep high humidity around the sample.

### Negative-stain Transmission electron microscopy

TEM images were obtained using a JEM-1400Flash (JEOL) operating at an accelerating voltage 120 kV. To observe the time course of self-assembly of TIR_MyD88_, TIR_MyD88_ solution (10∼300 μM) was incubated for 1 hour to 3 days at 4°C or 37°C. All the samples were left intact or diluted 10-fold in 20 mM HEPES pH 7.0, 50 mM NaCl, 5 mM DTT, loaded onto a carbon grid and stained with 1% uranyl acetate.

### High-speed AFM imaging

A high-speed atomic force microscope (HS-AFM) was equipped with a small cantilever (BL-AC10-DS-A2 (Olympus): spring constant, k = 0.1 N/m, resonance frequency, f = 400∼500 kHz in water) and was operated in tapping mode at room temperature ^2–4^. An amorphous carbon tip was grown on the top of each cantilever by electron-beam deposition with a scanning electron microscope (ERA-8000FE (Elionix)). The free oscillation amplitude was 1.4∼2 nm, and the typical set-point amplitude was 85% of the free oscillation amplitude. The imaging rate, scan size, and feedback parameters were optimized to enable visualization using a minimum tip force. A mica disk (1.5 mm in diameter) fixed by epoxy glue on a glass rod (1.5 mm in diameter and 2 mm in height) was used as a sample stage ^3^. After the sample stage was fixed by nail polish on the z-piezo of the HS-AFM scanner, the mica was freshly cleaved. Then, 2 μL of 10-200 μM TIR_MyD88_ was adsorbed on the substrates in AFM sample buffer containing 20 mM HEPES pH 7.0, 50 mM NaCl, and 10 mM DTT. After 10 min, the sample was rinsed with 20 μL of the sample buffer two times. The sample stage was filled with 63 μL of the sample buffer before starting measurement, and 7 μL of 50 μM GST-TIR_TLR2_, 50 μM TIR_TLR2_, 50 μM GST-TIR_TLR5_ or 50 μM GST was added to the sample stage while scanning.

### Processing of HS-AFM data

The HS-AFM image sequences were processed using Kodec 4.5.7.25 developed by the Kanazawa University WPI Nano Life Science Institute (WPI-NanoLSI) ^5^. The background height level was adjusted to zero after background leveling in all images. After taking images, we tracked a target molecule using two-dimensional (2D) correlation analysis to compensate for the slow drift of the sample stage position in the x- and y-directions. A 3 × 3 pixel-average filter was applied to each tracked image to reduce noise ^6^.

### Cryo-EM sample preparation

200 μM TIR_MyD88_ in 20 mM HEPES pH 7.0, 50 mM NaCl, 10 mM DTT was incubated for 3 days at 30 °C and 2.5 μL of the solution was applied onto a Quantifoil R 1.2/1.3 200 mesh Cu grid (Quantifoil) and plunge-frozen in liquid ethane by a Vitrobot Mark IV (FEI). The grids were glow-discharged with a 20 mA current for 30 s just before sample application. The parameters for plunge-freezing were as follows: blot time 2.5 s, humidity 100%, temperature 4 °C. After plunge-freezing, residual liquid ethane on the grid was thoroughly blotted off with filter paper and the grid were stored in liquid nitrogen.

### Image acquisition

Data acquisition was done by the CRYO ARM 300 (JEOL) operated at 300 kV installed with SerialEM ^7^ for automatic data collection and YoneoHole ^8^ for the hole alignment at Osaka University. The Ω-type incolumn energy filter was operated with a slit width of 20 eV for zero-loss imaging. The imaging parameters were: nominal magnification 50,000, defocus -0.5– - 2.0 µm, total dose 40 e^−^/Å^2^, exposure time 3.2 s, one image acquisition per a hole. Images were recorded with a K3 direct electron detector (Gatan) in CDS mode at a pixel size of 1.01 Å/pixel and each movie was fractionated into 40 frames.

### Image processing

A total of 5,325 images were collected. Image processing was performed with RELION 3.1 software ^9,10^. After motion correction with MotionCorr ^11^ and CTF estimation with Gctf ^12^, 4,930 images were selected for further image processing. The thinner cylindrical filaments were manually picked with e2helixboxer ^13^, after which 60,578 particles were extracted as overlapping boxes of 500 × 500 pixels with a step size of 66 Å along a filament. After 2D classification, 26,680 boxes were selected. Initial 3D models for searching a symmetry of the structure were created as assumed the circular symmetric bilayer using SPIDER ^14^, determined of C26 symmetry in 3D refinement. 3D reconstruction and refinement was performed with a helical refinement in Cryosparc software ^15^. Helical symmetry was determined to –9.2° twist/66.6 Å rise along the left-handed helix. Local and Global CTF refinement were performed. The final resolution reached 3.3 Å (Extended Data Fig. 2 and Supplementary Table 1). The structure determination of the thicker cylindrical fibers was conducted in a manner similar to the thinner ones described above. However, particle picking was performed using the filament tracer tool in Cryosparc, after which fibers of uniform diameter were manually selected and processed.

### Model building

The initial atomic model of TIR_MyD88_ were prepared from an atomic model of the crystal structure of the oligomeric state of the TIR_MyD88_ (PDB 7BEQ) ^16^. The model was fitted into the EM map using the ‘fit in map’ function of UCSF Chimera program ^17^ and was iteratively refined using COOT ^18^. Each of the final models was subjected to real space refinement in PHENIX ^19^.

### Structural analysis and visualization

PyMOL (version 2.2.3 Schrödinger, LLC), UCSF Chimera ^17^, UCSF ChimeaX ^20^ and PISA ^21^ were used to analyze and visualize the structures and to create the molecular graphics. The RMSD of the residues in the intra-strand interface was calculated using PyMOL based on the protein backbone atoms. The residues of the BB surface that interact with the EE surface, are P169, I172, Q173, Q176, I169 and S194-I207. The residues of the EE surface that interact with the BB surface, are K250, L252-P254, R269-C274, N278, C280, T281, W284, R288, L289, K291, A292 and L295.

## Data availability

The coordinates have been deposited in the Protein Data Bank (PDB) under accession code: TIR_MyD88_ oligomer forming stacked rings–8YYM and helical cylinder–8W8M, and the EM map has been deposited in the Electron Microscopy Data Bank (EMDB) (EMD-39676 and EMD-37355). All other data are available from the corresponding author upon reasonable request.

